# Satellite glial cells modulate proprioceptive neuron activity in dorsal root ganglia

**DOI:** 10.1101/2024.09.04.611156

**Authors:** Yasmine Rabah, Cendra Agulhon

## Abstract

Proprioception, the sense of body and limb position, is mediated by proprioceptors and is crucial for important motor functions such as standing and walking. Proprioceptor cell bodies reside within the peripheral dorsal root ganglia (DRG) and are tightly enveloped by satellite glial cells (SGCs). SCGs express a number of G_q_ protein-coupled receptors (G_q_ GPCRs), but their functional consequences on proprioceptor activity is unknown. Using a combination of chemogenetics, genetics, Ca^2+^ imaging, pharmacology, immunohistochemistry, and biochemistry, we provide evidence that SGC G_q_ GPCR signaling is sufficient to drive purinergic receptor-mediated Ca^2+^ responses in proprioceptor cell bodies. Our findings suggest a potential role for SGC G_q_ GPCR signaling in shaping proprioceptor information processing. Furthermore, this demonstration of SGC-induced proprioceptor activation has profound implications with SGC G_q_ GPCR signaling and purinergic receptors representing potential therapeutic targets for alleviating some proprioceptor and sensorimotor impairments associated with spinal muscular atrophy or Friedreich’s ataxia.

## Introduction

Proprioceptors innervate muscle spindles and tendons, and synapse onto spinal ventral horn lower motor neurons (MNs) to provide reflexive information about the length and contraction of muscles. They are crucial for coordinating the activity of MNs and skeletal muscles to achieve essential motor tasks, such as posture, locomotion, manoeuvring one’s way around obstacles or reacting rapidly to external perturbations^1,2^. When receiving sensory inputs at their peripheral nerve endings, DRG sensory neurons fire action potentials, eliciting the release of neurotransmitters from their cell bodies within DRGs^3^ as well as from their axonal terminals in the spinal cord. There is emerging evidence that transmitters released from sensory neuron cell bodies activate receptors at the surface of SGCs^4,5^ and that SGC-to-neuron purinergic communication takes place in DRG^11,12^. Both SGCs and somata of sensory neurons express a plethora of receptors, including G_q_ GPCR purinergic receptors^6–10^, however, only a few studies have examined the involvement of SGC G_q_ GPCR signaling in SGC-to-sensory neuron interactions and no study has specifically examined SGC-to-proprioceptor interactions.

Therefore, in this study, we asked whether selective activation of SGC G_q_ GPCR signaling is sufficient to elicit proprioceptor responses in DRGs. We further clarified whether purinergic receptors contributed to such proprioceptor responses. Our findings have identified, for the first time, a SGC-to-proprioceptor communication involving purinergic receptor-mediated Ca^2+^ activity in proprioceptors.

## Results

### Chemogenetic and genetic strategies for selectively activating SGC G_q_ GPCR signaling in DRGs

To define the role of SGC G_q_ GPCR signaling in proprioceptor activity, we used the GFAP-hM3Dq transgenic mouse model expressing a chemogenetic hM3Dq under the control of the glial GFAP promoter^13^. In DRGs, we found that hM3Dq is expressed in ∼88% of SGCs with no detectable expression in neurons, indicating that this mouse line represents a valuable model to active GPCR signaling selectively in the vast majority of SGCs (Fig. 1a, Fig. S1, Supplementary Movie 1 and Table 1).

**Figure 1.**
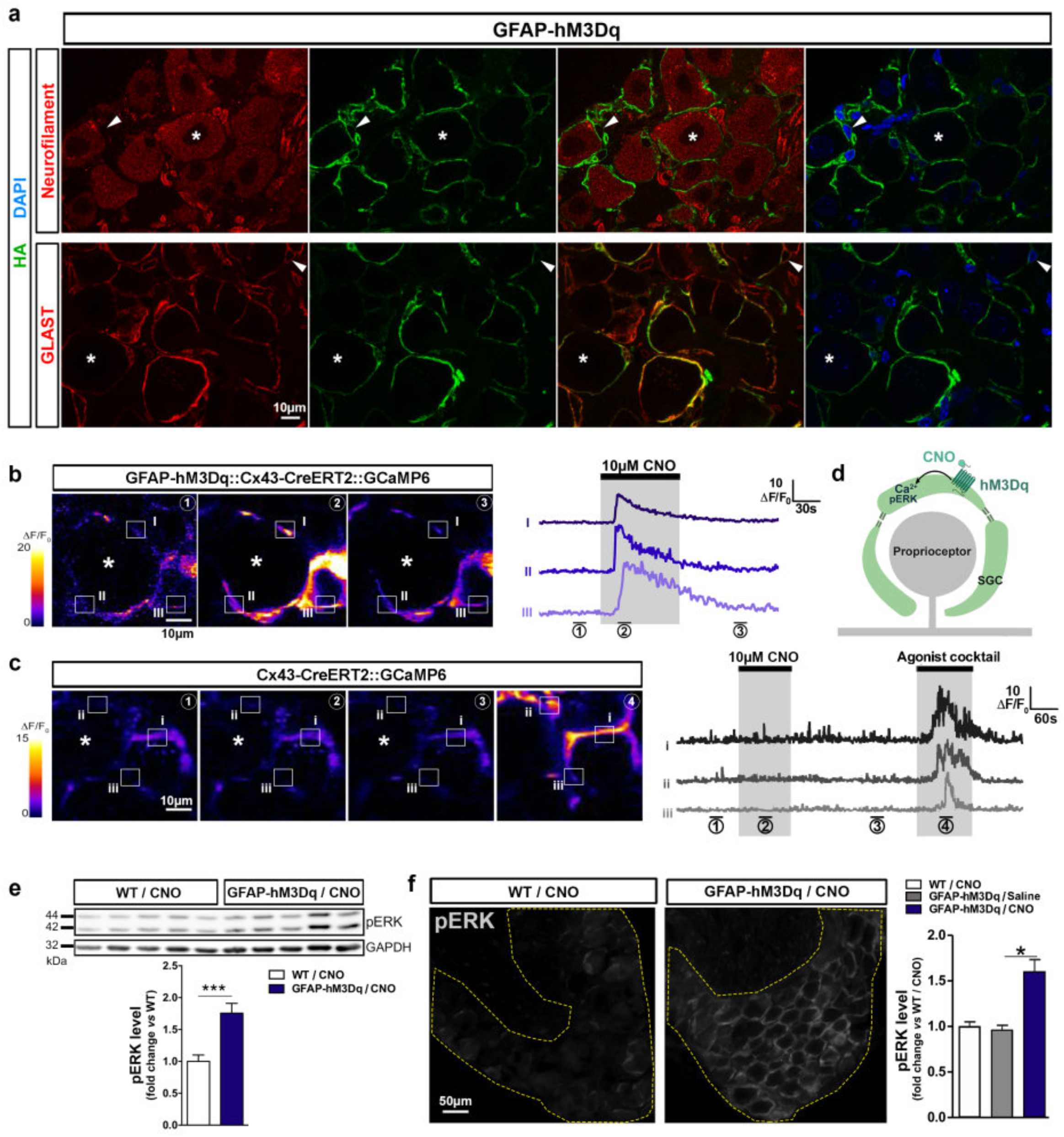
hM3Dq allows selective activation of G_q_ GPCR signaling cascades in DRG SGCs. **a**, Immunohistochemistry (IHC) in DRGs of GFAP-hM3Dq mice and confocal fluorescent images showing HA-tagged hM3Dq protein (green), sensory neurons labeled with neurofilament antibody (NF, red, top panel, asterisks) and SGCs labeled with GLAST antibody (red, bottom panel, arrowheads). Nuclei are stained with DAPI (blue). **b**, Two-photon Ca^2+^ imaging of *ex vivo* intact DRGs from GFAP-hM3Dq::Cx43-CreERT2::GCaMP6 mouse model. **Left panel** showing SGCs expressing GCaMP6f, outlined areas of interest (I-III) and Ca^2+^ increases during baseline ①, CNO application ②, and wash ③. Asterisk denotes neuronal cell soma. **Right panel** showing time course of CNO-mediated Ca^2+^ increases in SGCs (I-III). **c**, Calcium imaging of DRGs from Cx43-CreERT2::GCaMP6 mice. **Left panel** showing cells expressing GCaMP6f, outlined areas of interest (i-iii), and Ca^2+^ increases during baseline ①, CNO application ②, wash ③ and cocktail application ④. **Right panel** showing time course of Ca^2+^ traces in SGCs (i-iii). A cocktail of ligands to endogenous G_q_ GPCR (1µM adenosine, 50µM ATP-γS, 10µM carbachol, 50µM DHPG, 200µM glutamate, 10µM histamine) has been applied after CNO washing to ensure SCG viability. **d**, Schematic summarizing hM3Dq functionality in DRG SGCs. **e**,**f**, Western blot (WB) (**e**) and IHC (**f**) showing an increase of 76% (**e**) and 60% (**f**) of pERK expression level in GFAP-hM3Dq mice *vs.* control groups, 2min after *in vivo* CNO treatment (1mg/kg i.p.) (n = 9-10 animals/group for WB; n = 4 animals/group for IHC; two-tailed unpaired t-test for WB: *P* = 0.0006; Kruskal-Wallis test followed by Dunn’s multiple comparison test for IHC: *P* = 0.0231). GAPDH was used as a loading control in WB. \**P* < 0.05, \*\*\**P* < 0.001; error bars indicate mean ± SEM.

Then, to assess hM3Dq functionality in SGCs, we performed 2-photon Ca^2+^ imaging experiments in *ex vivo* intact DRGs from GFAP-hM3Dq::Cx43-CreERT2::GCaMP6 triple transgenic mice. These mice express hM3Dq and the genetically encoded Ca^2+^ indicator GCaMP6f (*fast*) selectively in SGCs (Fig. 1a and Fig. S2), enabling both hM3Dq-mediated Ca^2+^ elevations to be induced and detected selectively in SGCs. Indeed, bath application of 10 μM CNO elicited Ca^2+^ increases in ∼94% of SGCs, showing that, as expected, hM3Dq couples to G_q_ in these glial cells (amplitude: 16.62 ± 1.5 ΔF/F_0_; n = 119 SGCs from 9 DRGs and 3 mice; Fig. 1b,d and Supplementary Table 2). Importantly, no Ca^2+^ increase and no change in the frequency of spontaneous events were induced by CNO in DRGs from control Cx43-CreERT2::GCaMP6 double-transgenic mice^14^ (*i.e.* lacking hM3Dq expression in SGCs). However, a G_q_ GPCR agonist cocktail reliably triggered Ca^2+^ elevations in SGCs from these mice (n = 149 SGCs from 9 DRGs and 3 mice; Fig. 1c, Fig. S3 and Supplementary Table 2). Together, these data demonstrate that CNO has no non-specific effects in itself in our experimental conditions.

We next asked whether MAPK/ERK pathway was also activated downstream of hM3Dq stimulation in SGCs. Compared to control groups, DRGs from CNO-treated GFAP-hM3Dq mice (1mg/kg CNO intraperitoneal) exhibited a ∼76% increase in activated ERK1/2 (pERK) expression levels (Fig. 1e, Fig. S4 and Supplementary Table 3), which was found to be selective to SGCs and not to sensory neurons (Fig. 1f, Fig. S5 and Supplementary Table 3).

Together, these results validate the use of GFAP-hM3Dq mice for stimulating canonical G_q_ GPCR signaling cascades (*i.e.* Ca^2+^ and MAPK/ERK pathways) selectively in SGCs of DRGs.

### SGC G_q_ GPCR activation induces Ca^2+^ responses in proprioceptor cell bodies

To determine whether activating SGC G_q_ GPCR signaling is sufficient to modulate proprioceptor activity, we used *ex vivo* DRGs from another GFAP-hM3Dq::PV-Cre::GCaMP6 triple transgenic mouse model. In this model hM3Dq expression is under the control of GFAP promoter control (Fig. 1a), while GCaMP6f expression is under the control of parvalbumin (PV) promoter. In DRGs, parvalbumin is primarily found in proprioceptors. This model therefore enables the vast majority of SGCs to be activated by the application of CNO (Fig. 1b) while monitoring Ca^2+^ responses in ∼97% of proprioceptors expressing GCaMP6f (Fig. 2a,c,d; Fig. S6 and Supplementary Table 1). Notably, while proprioceptors exhibited no spontaneous Ca^2+^ transients in DRG preparations, 10µM CNO bath application elicited Ca^2+^ increases in a subpopulation of proprioceptors (amplitude: 1.26 ± 0.13 ΔF/F_0_; rise time: 23.6 ± 2.2 s; duration: 93.1 ± 3.5 s; Fig. 2a,c,d and Supplementary Movie 2 & Table 5).

**Figure 2.**
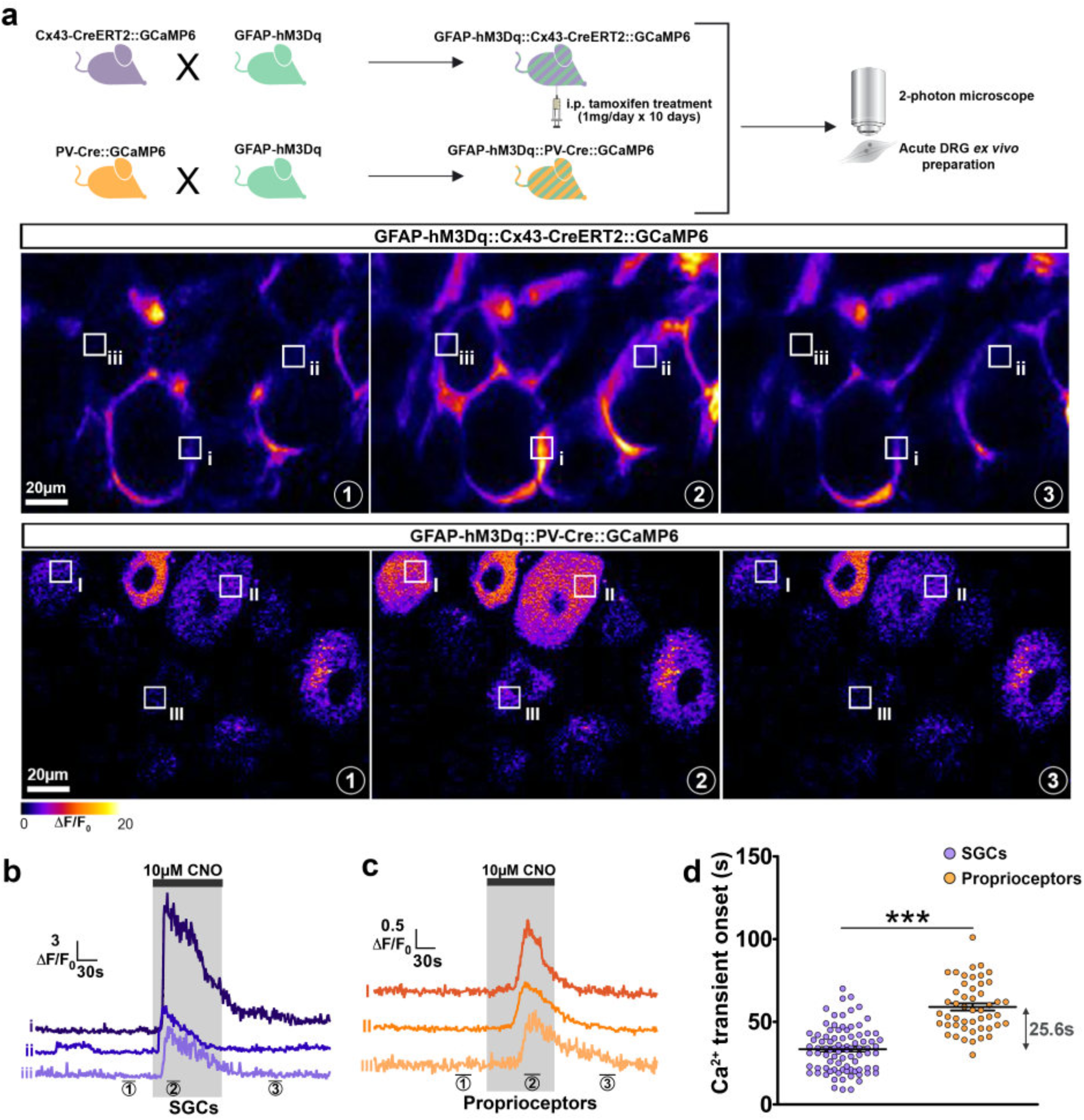
CNO/hM3Dq-mediated SGC activation induces proprioceptive neuron Ca^2+^ responses. **a**, Two-photon Ca^2+^ imaging of *ex vivo* DRGs from GFAP-hM3Dq::Cx43-CreERT2::GCaMP6 (top panel) and GFAP-hM3Dq::PV-Cre::GCaMP6 mouse lines (bottom panel) during baseline ①, CNO application ②, and wash ③, and outlined areas of interest (i-iii for SGCs and I-III for proprioceptors). **b**,**c**, Time course of CNO-mediated Ca^2+^ increases in SGCs from GFAP-hM3Dq::Cx43-CreERT2::GCaMP6 mice (**b**, i-iii) and proprioceptors from GFAP-hM3Dq::PV-Cre::GCaMP6 mice (**c**, I-III). **d**, Quantification of the onset of Ca^2+^ transients in SGCs and proprioceptors after bath application of CNO (n = 86 SGCs from 6 DRGs and 3 GFAP-hM3Dq::Cx43-CreERT2::GCaMP6 mice; n = 49 proprioceptors from 23 DRGs and 14 GFAP-hM3Dq::PV-Cre::GCaMP6 mice; Kruskal-Wallis test followed by Dunn’s multiple comparison test: *P*<0.0001; See Supplementary Table 4 for details).

Furthermore, upon 10 μM CNO application, we observed that the onset of these proprioceptor Ca^2+^ responses (in GFAP-hM3Dq::PV-Cre::GCaMP6 mice) occurred 25.6 ± 2.5 s after the onset of Ca^2+^ elevations elicited in SGCs (in GFAP-hM3Dq::Cx43-CreERT2::GCaMP6 mice) (2-tailed unpaired t-test, *P* = 0.0002; Fig. 2a-d and Supplementary Movie 3 & Table 4). This result was confirmed in DRGs obtained from a fourth mouse model expressing hM3Dq in SGCs as well as GCaMP6f in both SGCs and proprioceptors (GFAP-hM3Dq::Cx43-CreERT2::PV-Cre::GCaMP6 quadruple transgenic; Fig. 3a). Calcium invariably increased in SGCs first, followed by proprioceptors with a similar delay, demonstrating that SGC G_q_ GPCR activation drives (or alters) the activity of a subpopulation of proprioceptors (2-tailed unpaired t-test, *P* < 0.0001; Fig. 3b-e; Fig. S6 and Supplementary Movie 4 & Table 4). In support of this conclusion, CNO never induced Ca^2+^ elevations in proprioceptors of control PV-Cre::GCaMP6 double transgenic mice, demonstrating that proprioceptor Ca^2+^ responses are not due to a non-specific effect of CNO directly on neurons (Fig. S7 and Supplementary Table 2).

**Figure 3.**
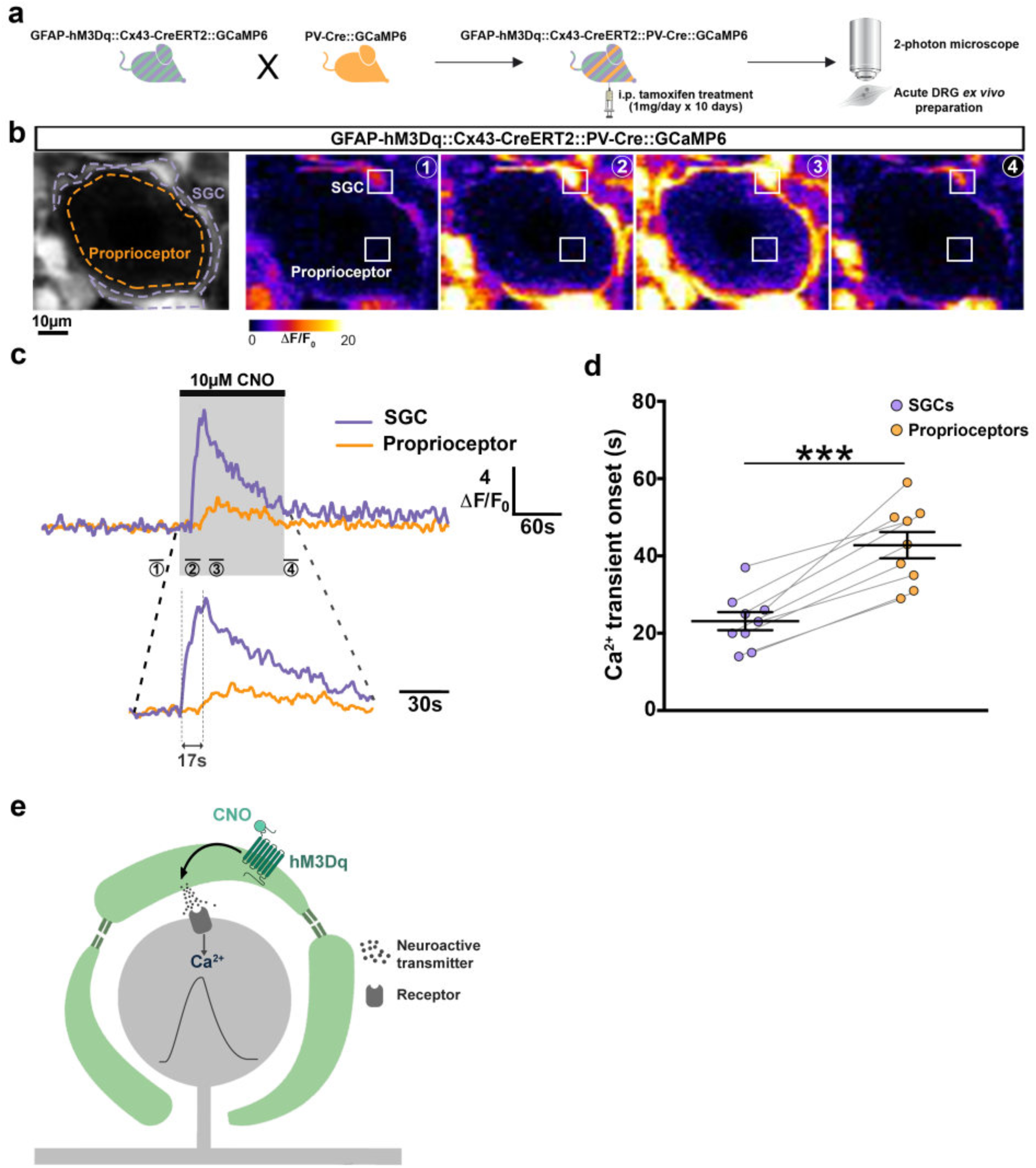
Direct demonstration that CNO/hM3Dq-induced activation of SGCs drives proprioceptor Ca^2+^ responses occurring ∼20 s later within same DRGs. **a,b,** Two-photon Ca^2+^ imaging of *ex vivo* DRGs from GFAP-hM3Dq::Cx43-CreERT2::PV-Cre::GCaMP6 quadruple transgenic mouse model, expressing hM3Dq in SGCs and GCaMP6f Ca^2+^ indicator in both SGCs and proprioceptors (**a**), during baseline ①, CNO application ②, and wash ③, and outlined areas of interest (SCG & proprioceptor) (**b**). The discrimination of SGCs and neurons is based on both morphological features and intensity of GCaMP6f brightness (left, black & white). **c**, Time course of CNO-mediated Ca^2+^ increases in SGC (purple) and neighboring proprioceptor (orange), and time axis enlargement. **d**, Quantification of the data showing that in pairs of SGCs and associated proprioreceptors, Ca^2+^ transients occuring in proprioreceptors invariably appears ∼20 s after CNO/hM3Dq-induced Ca^2+^ elevations in the surrounding SCGs (n=9 pairs each consisting of 1 SGC and 1 proprioceptor soma, n=5 DRGs, n=3 GFAP-hM3Dq::Cx43-CreERT2::PV-Cre::GCaMP6 mice. Two-tailed unpaired t-test: *P* = 0.0002. See Supplementary Table 4 for details). **e**, Schematic summarizing the hM3Dq-mediated and SGC-induced Ca^2+^ responses in neighboring proprioceptors. \*\*\**P* < 0.001; error bars indicate mean ± SEM.

### SGC-induced proprioceptor Ca^2+^ responses are mediated by purinergic receptors

We next aimed to determine which receptors were involved in this SGC-to-proprioceptor communication. Because functional ionotropic and metabotropic glutamate, GABA and ATP receptors are expressed at the plasma membrane of the soma of small-diameter DRG nociceptors^7,15,16^, we hypothesized that soma of large proprioceptive neurons may express similar receptors.

Similar 2-photon Ca^2+^ imaging experiments as previously described were performed using DRGs from GFAP-hM3Dq::PV-Cre::GCaMP6 mice. We found that concomitant bath application of 10 μM CNO and inhibitors of glutamatergic (AMPAR, NMDAR, groups I, II, III mGluRs), GABAergic (GABA_A_R, GABA_B_R) or purinergic (P2X_3_R and P2Y_1_R) receptors onto DRGs did not substantially depress the number of proprioceptors responding to CNO-induced SGC activation. Furthermore, proprioceptor Ca^2+^ response amplitude, rise time and duration were not significantly attenuated (Fig. 4a,b, Fig. S8 and Supplementary Table 5).

**Figure 4.**
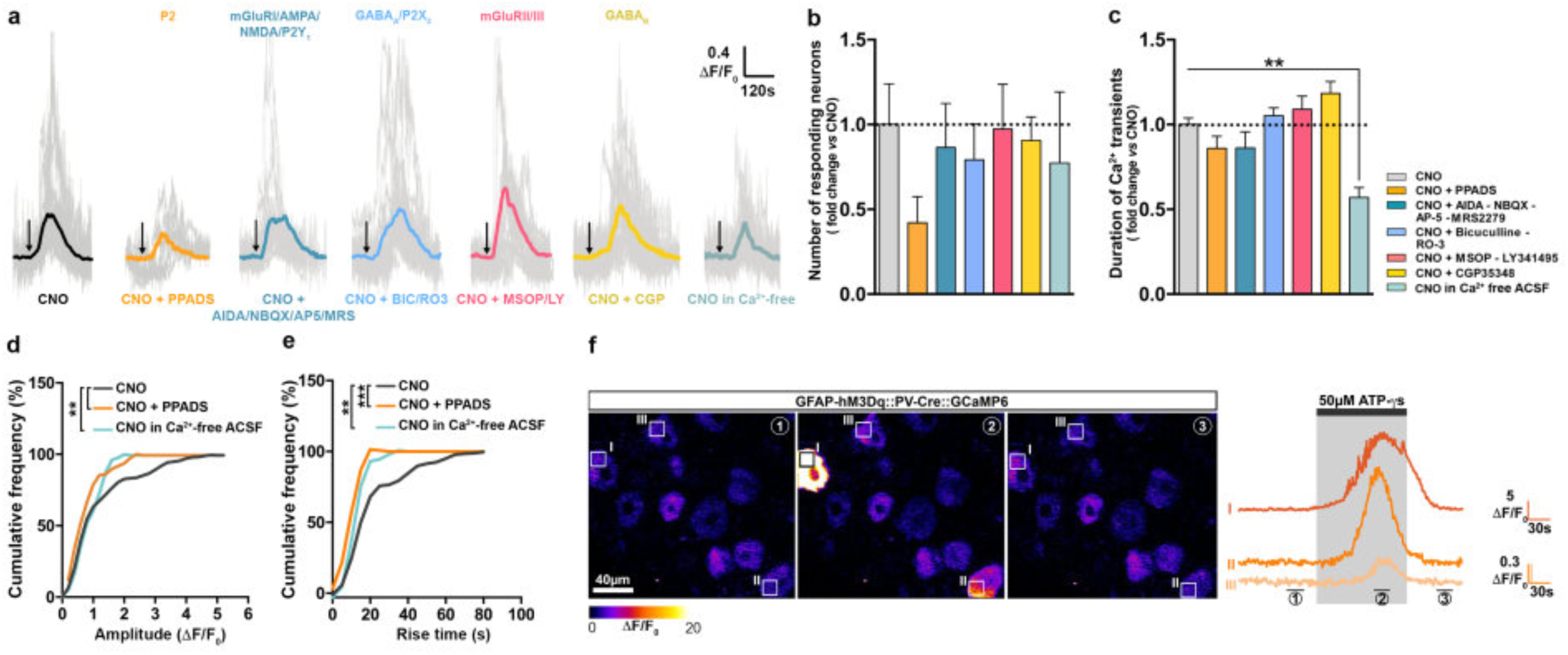
SGC G_q_ GPCR signaling induces Ca^2+^ responses in proprioceptors through purinergic receptors. **a**, Time course of CNO-induced Ca^2+^ increases in proprioceptors from GFAP-hM3Dq::PV-Cre::GCaMP6 mice following concomitant bath applications of CNO and antagonists to endogenous glutamatergic, GABAergic or purinergic receptors or to an application of CNO in extracellular Ca^2+^ free solution. **b**, Number of responding proprioceptors; the marked 57% decrease in the number of responsive proprioceptors when CNO is applied in presence of PPADS is not statistically significant (2-photon Ca^2+^ imaging data normalized to CNO condition; n = 49 DRGs from19 mice for CNO group and n = 16-19 DRGs from 5-6 mice for the other groups; Kruskal-Wallis test followed by Dunn’s multiple comparison test: *P* = 0.43; See Supplementary Table 5 for details). **c**, Duration of CNO/SGC-mediated Ca^2+^ increases in proprioceptors showing a significant decrease when CNO is applied in extracellular Ca^2+^ free solution (Kruskal-Wallis test followed by Dunn’s multiple comparison test: *P*<0.0001). **d**,**e**, Cumulative frequency distribution of proprioceptor Ca^2+^ response amplitudes and rise times (Kolmogorov-Smirnov tests; Amplitude: *P* = 0.0097 for CNO *vs.* “ CNO + PPADS” and *P* = 0.0014 for CNO *vs.* “CNO in Ca^2+^ free ACSF”; Rise time: *P* < 0.0001 for CNO *vs.* “ CNO + PPADS” and *P* = 0.0016 for CNO *vs.* “CNO in Ca^2+^ free ACSF”). **f**, Two-photon Ca^2+^ imaging of DRGs from GFAP-hM3Dq::PV-Cre::GCaMP6, outlined areas of interest (I-III, right panel), and time course of ATP-γS (50µM)-mediated Ca^2+^ increases in proprioceptors during baseline ①, ATP-γS application ②, and wash ③. \*\**P* < 0.01, \*\*\**P* < 0.001; error bars indicate mean ± SEM. Antagonist cocktails used and corresponding target receptors: [200 µM PPADS (P2R)];[100 μM AIDA (mGluR I) + 10 μM NBQX (AMPAR) + 50 μM AP5 (NMDAR) + 0.5 μM MRS2279 (P2Y_1_R)]; [50 μM Bicuculline (GABA_A_R) + 10 μM RO-3 (P2X_3_R)]; [100 μM MSOP (mGluR II) + 1 μM LY341495 (mGluR III)] ; and [100 μM CGP35348 (GABA_B_R)]. \*\**P* < 0.01, \*\*\**P* < 0.001; error bars indicate mean ± SEM.

However, co-applying 10 μM CNO with the broad-spectrum purinergic receptor antagonist PPADS (100 μM) reduced proprioceptor Ca^2+^ increase amplitude and rise time by ∼34% and ∼49% as shown in the left shifted cumulative probability distributions (0.83 ± 0.16 ΔF/F_0_; 12.1 ± 1.3 s; Kolmogorov-Smirnov test: *P* = 0.0348, *P* = 0.0007; Fig. 4d,e), with no change in Ca^2+^ response duration (Fig. 4c, Fig. S8 and Supplementary Table 5). A decrease (∼57% drop) in the total number of responsive proprioceptors was also observed (Fig. 4b). Taken together, these results suggest that proprioceptor purinergic receptors - other than P2X_3_R and P2Y_1_R previously reported in nociceptors^11,12^ - and presumably ATP released by SGCs, mediate Ca^2+^ increases in proprioceptors.

We therefore examined whether proprioceptor cell bodies could respond to ATP through functional purinergic receptors. Adding 50 μM non-hydrolyzable ATPγS on DRGs from PV-Cre::GCaMP6 mice elicited Ca^2+^ responses in ∼33% of proprioceptors as compared to CNO application (Fig. 4f and Supplementary Table 6). Thus, a subpopulation of proprioceptors express purinergic receptors at their cell body plasma membrane. The fact that nonhydrolyzable ATPγS did not activate as many proprioceptors as CNO did, may suggest that another transmitter (*e.g*. ADP *via* ATP hydrolysis) is required for the full effectiveness in SGC-to-proprioceptor activation. In agreement with this hypothesis, 30 μM nonhydrolyzable ADPβS induced Ca^2+^ responses in ∼46% of proprioceptors as compared to CNO (Fig. S9 and Supplementary Table 6).

### Proprioceptor Ca^2+^ responses require both extracellular and intracellular Ca^2+^

We next investigated the possible contribution of extracellular Ca^2+^ to the CNO-induced/SGC-mediated proprioceptor responses. To this end, a Ca^2+^ free extracellular solution was used to perform 2-photon imaging experiments, always using DRGs from GFAP-hM3Dq::PV-Cre::GCaMP6. We observed that the number of responsive proprioceptors was not appreciably modified upon 10 μM CNO application in an Ca^2+^ free extracellular solution (Fig. 4a,b), indicating that extracellular Ca^2+^ is not required for the induction of proprioceptor responses. However, the response amplitude, rise time and duration were markedly attenuated by 25%, 38% and 43%, respectively, suggesting that extracellular Ca^2+^ is required for reaching the largest proprioceptor responses (0.94 ± 0.12 ΔF/F_0_; 14.6 ± 1.8 s; 53.0 ± 5.4 s; Kolmogorov-Smirnov test: *P* = 0.0047, *P* = 0.007 and Kruskal Wallis test: *P* < 0.0001; Figure 4a,c-e, Fig. S8 and Supplementary Table 5).

The contribution of extracellular Ca^2+^ was further substantiated by the observation of a ∼41s delay in the proprioceptor response onset compared to the response onset obtained in regular extracellular solution condition (*i.e.* containing Ca^2+^) (Fig. S10 and Supplementary Table 4).

Overall, our data are consistent with a model in which activation of both ionotropic P2XR (mediating Ca^2+^ entry from extracellular space) and metabotropic P2YR (mediating Ca^2+^ release from internal stores) contribute to the full proprioceptor cytosolic Ca^2+^ responses. The P2XR-mediated Ca^2+^ increase preceding a subsequent delayed metabotropic P2YR-mediated Ca^2+^ increase. Such type of synergistic interactions between P2XR and P2XR has been previously described in marrow megakaryocytes and blood platelets^17^.

## Discussion

The primary goal of this study was to address the role of SGC G_q_ GPCR signaling in proprioceptor activity. Collectively, our findings reveal a new mechanism of interaction between SGCs and proprioceptors involving, at least partially, purinergic P2XR and P2YR as well as ATP and/or ADP transmitters.

Although scarce, studies have suggested that ATP release from SGCs results in the co-activation of neuronal P2Y_1_R and P2X_3_R and Ca^2+^ rises in nociceptive neurons, with P2Y_1_R exerting an inhibitory action on the function of P2X_3_R^4,5,10^. However, our data show that these purinergic receptors are not involved in SGC-mediated proprioceptor Ca^2+^ responses. Indeed, selective antagonists of these two receptors do not prevent or alter Ca^2+^ responses in proprioceptors (Fig. 4). Instead, our results suggest that SGCs exert control on proprioceptor activity *via* other purinergic receptors (blocked by pan PPADS broad-spectrum purinergic antagonist). Thus, the purinergic system appears to be a conserved mechanism for SGC communication with both nociceptors and proprioceptors, but to involve different purinergic signaling modalities depending on the type of sensory neurons. To the best of our knowledge, differential communication between SGCs and different types of sensory neurons has never been documented before. Nevertheless, it represents a form of communication whereby SGCs could discern specific populations of sensory neurons and induce distinct control/alteration of the Ca^2+^ homeostasis of these neurons depending on the involved sensory modality or the physiopathological state.

Additionally, the partial reduction of SGC-induced proprioceptor Ca^2+^ responses in the presence of PPADs can be explained by the fact that PPADS is a general inhibitor of purinergic receptors that does not inhibit all purinergic receptors. Therefore, it remains possible that residual proprioceptor Ca^2+^ activity is due to the lack of inhibition of certain purinergic receptors, or possibly to the indirect opening of plasma membrane Ca^2+^ channels downstream of purinergic receptor activation. Elucidating these questions and mechanisms is beyond the scope of the present study. It will require using a combination of available purinergic receptor knockout animals and purinergic receptor and Ca^2+^ channel antagonists, which represent a considerable endeavor.

Clear proof of expression of functional purinergic receptors on proprioceptor cell bodies has not yet been described. Our data showing that non-hydrolyzable ATPγS and ADPβS induce Ca^2+^ elevations although additional indirect scenario involving another cell type cannot be totally ruled out. Furthermore, a recent transcriptional profiling study has uncovered that mRNAs of purinergic ionotropic P2X_5_R and P2X_6_R as well as metabotropic P2Y_14_R are enriched in proprioceptors^18^. Although the presence of these transcripts does not indicate that the corresponding receptors are expressed at the level of proprioceptor soma, P2X_5_R, P2X_6_R and P2Y_14_R represent interesting candidates potentially involved in SGC-to-proprioceptor communication. As mentioned above, further studies are necessary to address this hypothesis.

Not only proprioceptors, but also low-threshold mechanoreceptors, express parvalbumin. However, our PV-Cre::GCaMP6 mouse model (in which parvalbumin promoter controls GCaMP6f expression) exhibits the presence of the Ca^2+^ indicator primarily in proprioceptors (Fig. S6), suggesting that it is a suitable model to detect intracellular Ca^2+^ homeostasis changes specifically in proprioceptors. Yet, the possibility that a few low-threshold mechanoreceptors also express GCaMP6f and their activity is included in our data sets cannot be excluded.

Finally, irrespective of mechanism(s), our results raise the possibility that SGC G_q_ GPCR signaling-induced disruption of activity in a subpopulation of proprioceptors might be sufficient to modulate some type of proprioceptive information processing within DRGs and produce changes in sensorimotor behavior. In conclusion, our study is relevant to proprioceptive impairments as well as sensorimotor deficits and ataxia associated with spinal muscular atrophy or Friedreich’s ataxia, respectively^19–21^

## Materials and Methods

### Animals

Experiments were conducted on 2- to 3-month old male and female mice from the C57BL/6N background. Mice were grouped housed (5 mice/cage) and fed *ad libitum*. Illumination was controlled automatically with a 12/12h light-dark schedule. All experiments were conducted during the dark phase. Wildtype (WT) littermates were used as controls in experiments involving transgenic mice. The following mouse lines were used and/or generated (the mouse lines used in experiments appear in italic; Supplementary Table 7): (1) PV-Cre^22^ (Jackson Laboratories, stock #017320) and Cx43-CreERT2^23^ mice were crossed with CAG-lox-STOP-lox-GCaMP6^24^ (Jackson Laboratories, stock #024105) in order to obtain two new double transgenic mouse lines that we called *PV-Cre::GCaMP6* and *Cx43- CreERT2::GCaMP6*, respectively. These two lines were used for running control Ca^2+^ imaging experiments to test the internees of CNO (e.g. to test whether or not CNO in itself evokes Ca^2+^ elevations were used for Western blot, immunohistochemical and behavioral experiments. These mice were crossed with both *PV-Cre::GCaMP6* and *Cx43-CreERT2::GCaMP6*^14^ transgenic mice in order to obtain two new triple transgenic mouse line that we named *GFAP-hM3Dq::PV-Cre::GCaMP6* and *GFAP- hM3Dq::Cx43-CreERT2::GCaMP6*, respectively. These two triple transgenic mouse lines were used for Ca^2+^ imaging experiments. Finally, we crossed *GFAP-hM3Dq::Cx43-CreERT2::GCaMP6* with *PV-Cre::GCaMP6* in order to obtain a new quadruple transgenic mouse line that we called *GFAP- hM3Dq::Cx43-CreERT2::PV-Cre::GCaMP6*. To induce GCaMP6f expression in *Cx43- CreERT2::GCaMP6*^14^ and in *GFAP-hM3Dq::*Cx43-CreERT2::GCaMP6, mice were treated i.p. with tamoxifen (1mg/day, Sigma) diluted in corn oil (Sigma) during 10 days and used 15 days after the first day of treatment. Animal care and procedures were carried out according to the guidelines set out in the European Community Council Directives.

### Immunohistochemistry

Animals were sacrificed 2min after treatment (CNO 1mg/kg or 0.9% NaCl, i.p.) and DRGs were immediately harvested and drop fixed in 4% paraformaldehyde for 4h prior to cryoprotection in 0.02M PBS containing 20% sucrose overnight at 4°C. For other immunohistochemistry experiments (e.g. transgenic mice characterization), animals were transcardiacally perfused with 4% paraformaldehyde under ketamine/xylazine (100mg/kg / 10mg/kg respectively, i.p.) anesthesia. Lumbar L3, L4 and L5 DRGs were harvested, postfixed 2 h in 4% paraformaldehyde, and cryoprotected in 0.02M PBS containing 20% sucrose overnight at 4°C. Then, tissues were frozen in optimal cutting temperature compound (OCT). Fourteen µm sections were cut using a cryostat (Leica), mounted on Superfrost glass slides and stored at −80°C. The day of the experiment, sections were washed two times for 15 min each in 0.02M PBS. Sections were incubated overnight in 0.02M PBS (pH 7.4) containing 0.3% Triton X100, 0.02% sodium azide and primary antibodies (Supplementary Table 8) at room temperature in a humid chamber. The following day, sections were washed 3 times in 0.02M PBS, and incubated for 2h at room temperature with secondary antibodies diluted in 0.02M PBS (pH 7.4) containing 0.3% Triton X100 and 0.02% sodium azide. Then, sections were washed 3 times for 15min in 0.02M PBS and mounted between slide and coverslip with Vectashield medium containing DAPI (Vector Laboratories). Negative controls, i.e. slices incubated with secondary antibodies only, were used to set criteria (gain, exposure time) for image acquisition for each experiment. Images acquisition was performed with an Axio Observer Z1 epifluorescence Zeiss microscope, an ORCA Flash 2.8 million pixel camera, a PlanNeoFluar 20x/0.5NA objective, a LED COLIBRI 2 light source with 4 narrowband LED (365nm, 470nm,590nm, 625nm), as well as filters [DAPI (49), eGFP (38HE), Cy3 (43HE) and Cy5 (50)]. A Zeiss LSM710 confocal microscope with a Plan-apochromat 63x oil immersion objective (umerical aperture of 1.4) was also used. The same image acquisition settings were used for the negative control (Zeiss). Cell counting and mean grey value measurements were performed using ImageJ software software from the National Institute of Health (USA). Because it was difficult to discriminate individual SGCs within all SGCs surrounding a single neuronal cell body, “rings” surrounding neuronal cell bodies were quantified. For confocal imaging of GFAP-hM3Dq DRG sections, 12µm thick z-stacks were acquired with 0.3µm steps (40 z-stacks in total). 3D visualization was done using the 3D viewer plugin of Fiji. Raw data were analyzed and quantified.

### Western Blots

Animals were sacrificed 2min, 30min and 4h after treatment (CNO 1mg/kg i.p.) and L3, L4, L5 DRGs were dissected, frozen and stored at −80°C. The tissues were homogenized in 150μL RIPA buffer (50mM Tris pH 8, 150mM NaCl, 1% NP-40, 0,5% deoxycholate, 0,1% SDS, 1 mM sodium orthovanadate) with 1X protease inhibitor cocktail cOmplete (Roche) and 1X Halt phosphatase inhibitor cocktail (Thermo Scientific). Tissue lysate was obtained using a Bioruptor sonication system (Diagenode) and then centrifugated at 2000 RPM for 5min. Supernatant was kept and protein concentration was determined using a BCA assay (Bio-Rad). Aliquots of 25µg of protein for each mouse were deposited and run on 10% acrylamide SDS-page and transferred to nitrocellulose membranes. Membranes were cut (according to protein weight), and the pieces of membranes were then saturated in TBS-Tween (TBS-T) containing 5% fat-free milk for 30min. Membranes were then incubated overnight at 4°C in TBS-T containing primary antibodies (Supplementary Table 8). The day after, membranes were washed in TBS-T and then incubated with horse radish peroxidase (HRP)-conjugated secondary antibodies diluted in TBS-T (Supplementary Table 8) for 1h30 at room temperature. Clarity ECL chemiluminescence detection (Biorad) and ImageQuant LAS4000 (GE Healthcare Life Sciences) were used to reveal and visualize the proteins. The average exposure time was 30s and images were taken with 1s increments. The images obtained before saturation of the signal were used for quantification. The mean gray values corresponding to the signal were measured using ImageJ software. Western blot experiments were replicated 4 times to minimize the technique variability and the 4 mean grey values obtained per sample were then averaged. Then each average value was normalized to the values obtained for the control group. Furthermore, to average away any possible position effect, the order of the deposits was different in each replicate.

### Two photon Ca^2+^ imaging

Acute intact DRG preparations were prepared from GFAP-hM3Dq::PV-Cre::GCaMP6, GFAP-hM3Dq::Cx43CreERT2::GCaMP6, and GFAP-hM3Dq::PV-Cre::Cx43-CreERT2::GCaMP6 mouse lines. PV-Cre::GCaMP6 and Cx43CreERT2::GCaMP6 mice were also used to perform prior control experiments to test the inertness of CNO. Vertebras and dura mater were removed and L4 and L5 were cold (slushy) incubation ACSF solution containing (in mM): 95 NaCl, 1.8 KCl, 1.2 KH_2_PO_4_, 0.5 CaCl_2_, 7 MgSO_4_, 26 NaHCO_3_, 15 glucose, and 50 sucrose, oxygenated with 95% O_2_ and 5% CO_2_. DRGs were harvested and incubated at 35°C for 30min in incubation solution and then left to recover for 1h30 at room temperature. A single DRG was placed in the recording chamber of a custom-built two-photon laser-scanning microscope with a 20x water immersion objective (x20/0.95w XLMPlanFluor, Olympus). GCaMP6f was excited at 920nm with a Ti:Sapphire laser (Mai Tai HP; Spectra-Physics). DRGs were continuously superfused with oxygenated recording solution identical to the incubation solution except for the following (in mM): 127 NaCl; 2.4 CaCl_2_; 1.3 MgSO_4_; and 0 sucrose at a rate of 4 ml/min. For experiments in Ca^2+^-free ACSF, the recording solution was identical to incubation solution except the following (in mM) : 127 NaCl; 0 CaCl_2_; 1.3 MgSO_4_; and 0 sucrose. Image acquisition was performed at a rate of 1 image/second (1Hz). Drugs applied are detailed in Supplementary Table 8. To determine the viability of proprioceptors GFAP-hM3Dq::PV-Cre::GCaMP6 mice, KCl (50mM) was applied as a positive control at the end of every experiment (Supplementary Movie 5); the number of proprioceptors responding to KCl represented the total number of proprioceptors indicated in Supplementary Tables 4, 5, 6. To determine the viability of SGCs in GFAP-hM3Dq::Cx43CreERT2::GCaMP6 and Cx43CreERT2::GCaMP6 mice, we considered SGCs responding to CNO and/or agonist cocktail to endogenous G_q_ GPCRs (50 μM DHPG, 10 μM histamine, 10 μM carbachol, 50 μM ATP-γS, 1 μM adenosine, 200 μM glutamate) as the total number of alive SGCs. Agonist cocktail was systematically applied at the end of every experiment. ImageJ and Metamorph (Molecular Devices) softwares were used to analyze the data. Regions of interest were determined on GCaMP6f-expressing cells and the relative changes in fluorescence (ΔF/F_0_) were calculated as the ratio of the fluorescence intensity to that recorded before any drug application. In GFAP-hM3Dq::Cx43-CreERT2::PV-Cre::GCaMP6 mice, in which GCaMP6f expression is driven both in SGCs and proprioceptors, the two cell types were discriminated by their distinct morphological features and GCaMP6f brightness differences (indeed SGCs expressed more GCaMP6f than proprioceptors and thus were brighter than proprioceptors in basal conditions). Positive responses were defined as those that exceeded 3 standard deviations (SD) above the baseline level. To determine the onset of Ca^2+^ responses, the time for the drug-containing ACSF to reach the recording chamber through the tubing was subtracted. Rise times were calculated as the 10-90% rise time. The designs of the different Ca^2+^ imaging experiments are presented in Supplementary Fig.11.

### Statistics

Data were acquired and analyzed blind of genotype and treatment. Data are shown as mean ± S.E.M. *n* values correspond to the number of cells, DGR, or mice. All statistical tests were performed after verification of normal data distribution using D’Agostino and Pearson omnibus normality test and, when applicable, equality of variances with F-test (Fisher). If normality assumptions were not met, we used Dunn’s multiple comparison test for analyzing more than 2 groups). If normality was met, we used parametric tests (two-tailed Student t-test for analyzing 2 groups and one-way ANOVA followed by Tukey post-hoc for analyzing more than 2 groups). Kolmogorov-Smirnov test was used to analyze cumulative distributions of pairs of groups. All statistical tests were performed using GraphPad Prism 6. No statistical methods were used to pre-determine sample sizes.

## Author contributions

C.A. and Y.R. designed and interpreted experiments. Y.R. performed experiments and data analysis.

Y.R. wrote the first draft of the manuscript. C.A. edited and wrote the manuscript. C.A. conceived and supervised the project.

## Acknowledgments

We gratefully acknowledge C. Steinhäuser for providing the Cx43-CreERT2 mouse line; S. Antoine and S. Guinoiseau for animal care; J.M. Andrieu, F. Charbonnier, B. Delhomme, P. Djian, C. Levenes, C. Meunier and M. Oheim for sharing pieces of equipment and laboratory spaces; O. Biondi and E. Schmidt for advice and support on image acquisition; A. Bessis and S. Dieudonné for valuable discussions and feedback; T. Fiacco and J. Stinnakre for critical reading and I. Melnychuk for editing the manuscript; and both the imaging and mouse core facilities, which are supported and funded by CNRS, INSERM and Université Paris Cité.

## Supplementary information

**Supplementary Figure 1 (related to Fig. 1a).**
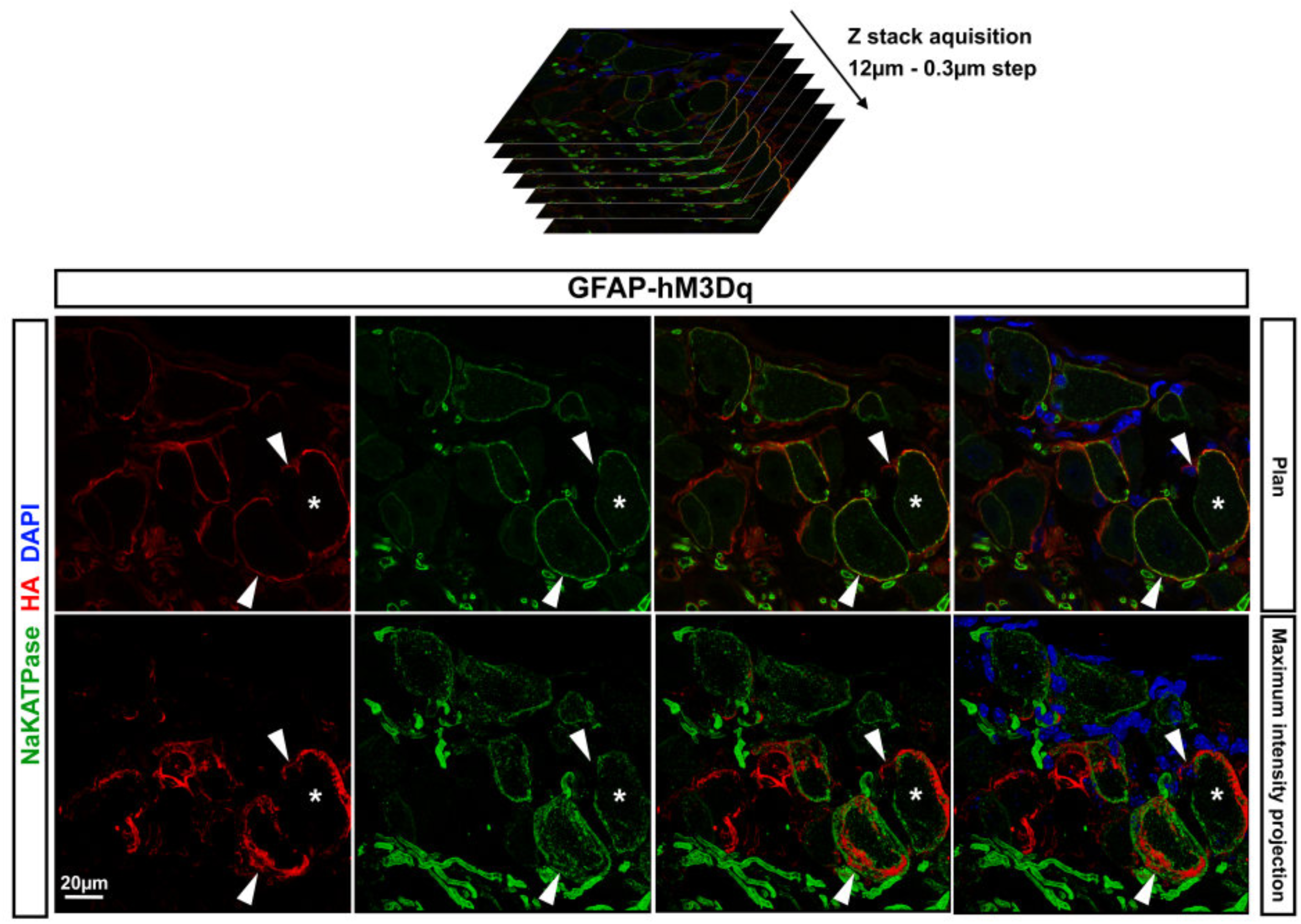
Cellular characterization of GFAP-hM3Dq mouse line. Immunohistochemical visualization of HA-tagged hM3Dq protein (HA, red) and Na/K-ATPase neuronal plasma membrane marker (green) in DRGs from GFAP-hM3Dq mouse. **Top panel**, Single plan imaged with a confocal microscope. **Bottom panel**, Since the distance between SGC and neuronal plasma membranes is about 20nm, i.e. beyond confocal microscope resolution, z-optical sections were stacked to explore possible colocalization between HA and Na/K-ATPase staining: 40 z-optical sections, 12 µm each, with a step of 0.3 µm were stacked to obtain a maximum intensity projection. Areas expressing HA and Na/K-ATPase are different, showing that HA-tagged hM3Dq is not expressed at the plasma membrane of sensory neurons. Nuclei of HA-expressing SGCs (stained in blue with DAPI, arrowheads) appear small with a bright DAPI labeling. This is different from neuronal nuclei, which are large with a dim DAPI staining (asterisks).

**Supplementary Figure 2 (related to Fig. 1b,c).**
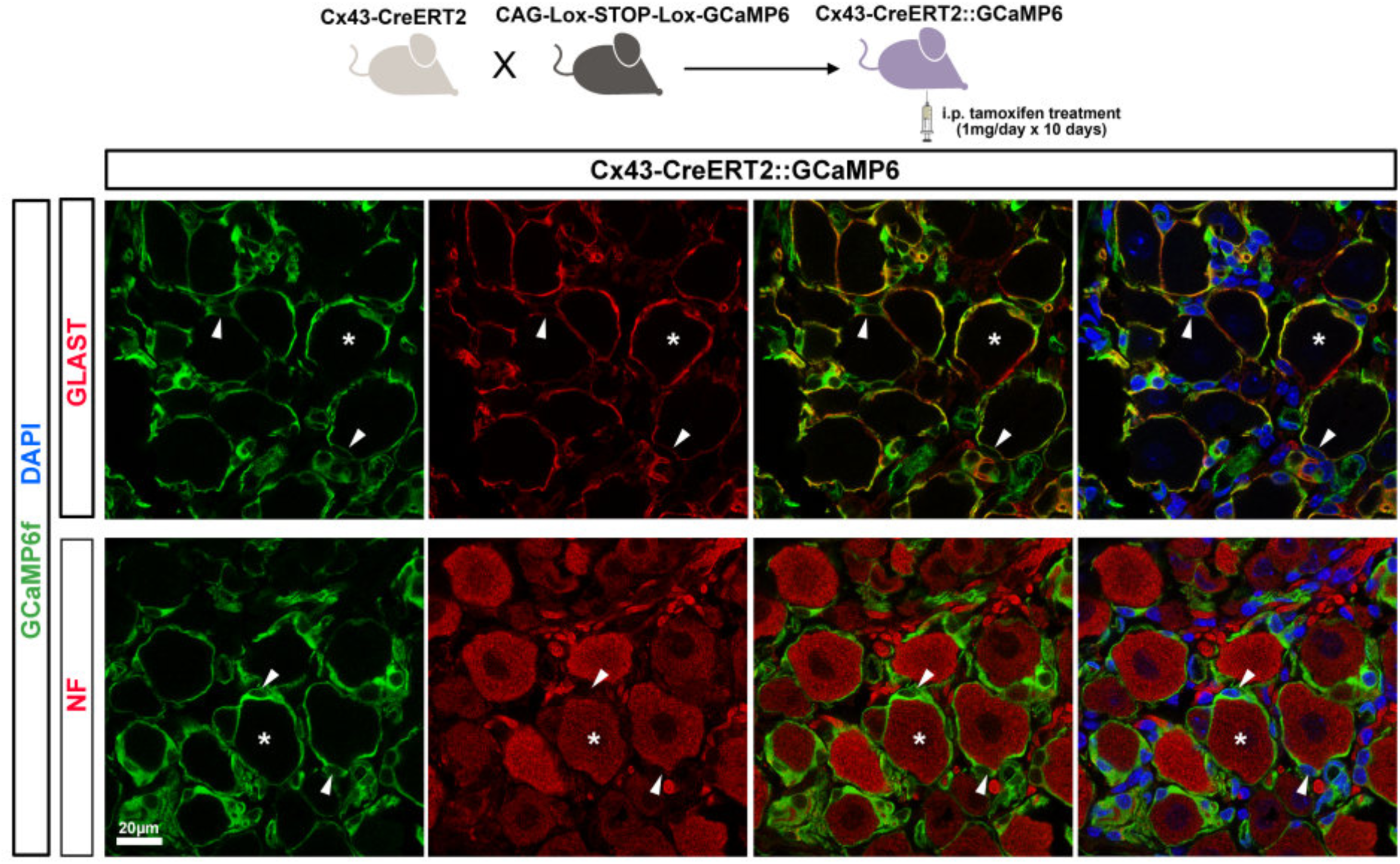
Cellular characterization of Cx43-CreERT2::GCaMP6 double transgenic mouse line. **Top panel**, Confocal images of immunohistochemical experiments showing GCaMP6f (green) overlapping with SGC GLAST marker (red). **Bottom panel**, GCaMP6f was not detected in neurofilament-expressing cell bodies of sensory neurons (red). Arrowheads and asterisks denote GLAST-expressing SGCs and neurofilament-expressing neurons, respectively. SGC and neuronal cell bodies are stained with DAPI (blue).

**Supplementary Figure 3 (related to Fig. 1b,c).**
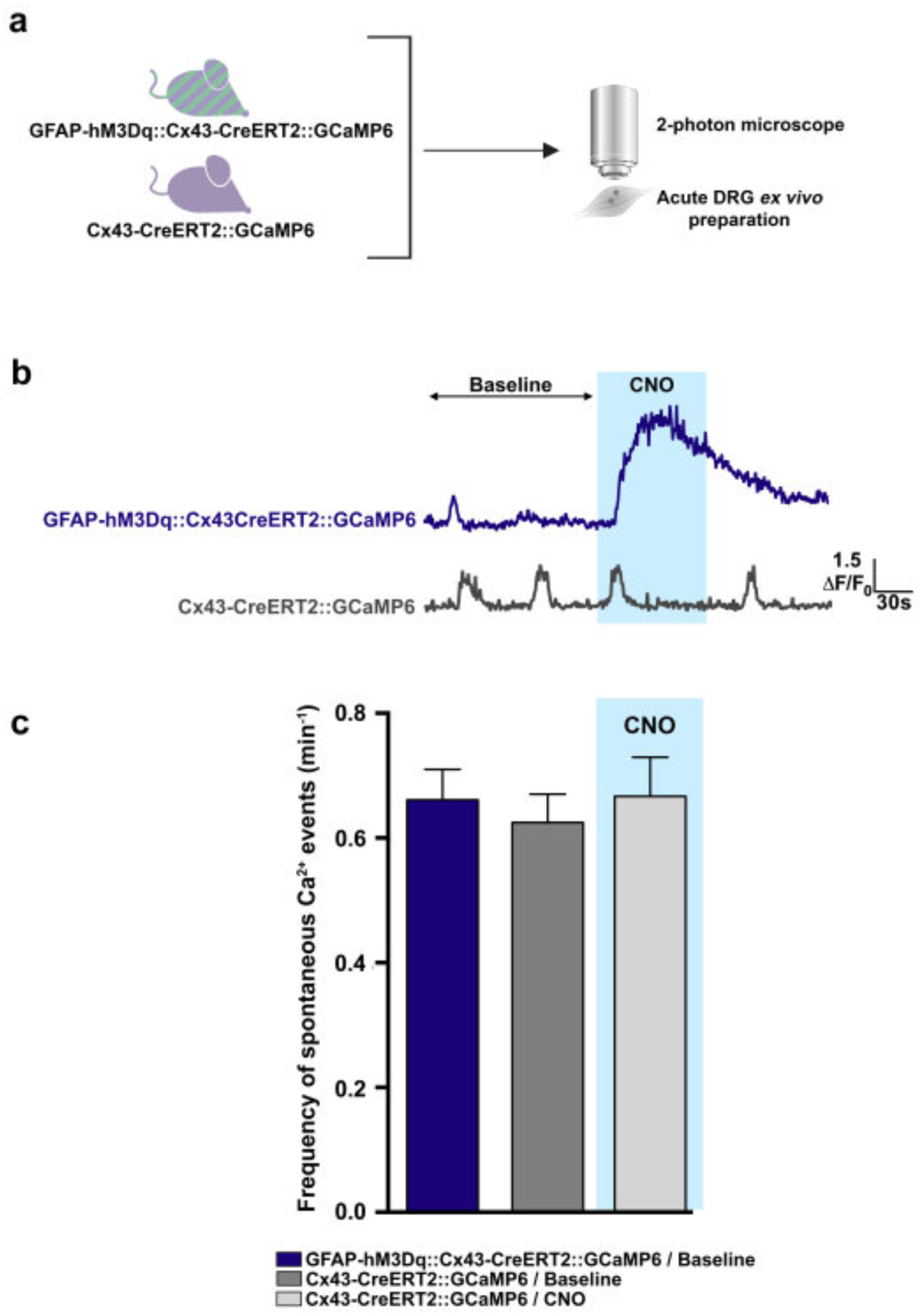
Inertness of CNO on the frequency of spontaneous Ca^2+^ events in SGCs and absence of hM3Dq receptor constitutive activity. In Cx43-CreERT2::GCaMP6 double transgenic mouse line, spontaneous Ca^2+^ elevations occurred in ∼12% of SGCs. **a**, Schematic of experimental design. **b**, Calcium traces showing the pattern of expression of spontaneous Ca^2+^ events compared to CNO-evoked Ca^2+^ elevations in SGCs. **c**, Bath application of 10 μM CNO to *ex vivo* DRGs from Cx43-CreERT2::GCaMP6 mice does not modulate (increase or decrease) the frequency of spontaneous Ca^2+^ elevations compared to baseline (2min before CNO application). This result indicates that CNO has no non-specific effect in itself in agreement with the data reported in Fig. 1c. Both GFAP-hM3Dq::Cx43-CreERT2::GCaMP6 and Cx43-CreERT2::GCaMP6 control mice, exhibit similar frequency in SGC spontaneous Ca^2+^ elevations during baseline, suggesting that hM3Dq receptor has no constitutive activity in itself. Data quantification: Cx43-CreERT2::GCaMP6 mice, during CNO application: 0.67 event/min; Cx43-CreERT2::GCaMP6 mice, during baseline: 0.63 events/min; GFAP-hM3Dq::Cx43-CreERT2::GCaMP6 mice, during baseline: 0.66 event/min. n=24 cells from Cx43-CreERT2::GCaMP6 mice, n=31 cells from GFAP-hM3Dq::Cx43-CreERT2::GCaMP6 mice. Kruskal-Wallis test followed by Dunn’s Multiple Comparison Test: *P*= 0.8503.

**Supplementary Figure 4 (related to Fig. 1e).**
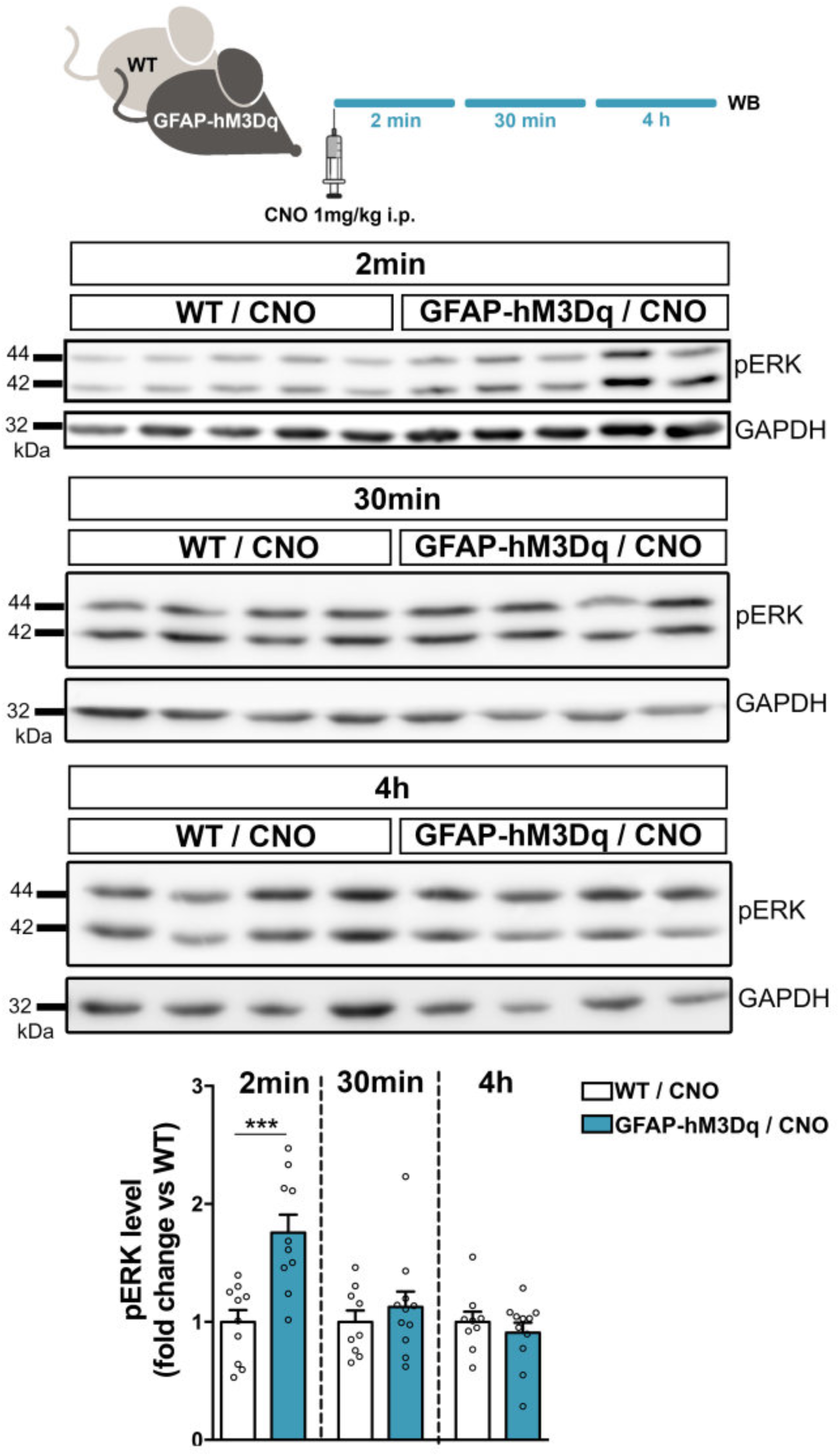
Western blot showing the timeline of MAPK/ERK pathway activation downstream of hM3Dq stimulation in DRGs *in vivo*. **Top panel**, Experimental design. GFAP-hM3Dq and WT littermate control mice were treated intraperitoneally with 1mg/kg CNO, sacrificed 2min, 30min or 4h after treatment, and the L3, L4 and L5 DRGs (in total 6 DRGs per mouse) were dissected and processed for Western blot experiments. Note that the average time intervals between the dissection and freezing procedure of the 1^st^ DRG and the 6^th^ DRG were ∼1min 30s and ∼3min 40s, respectively. **Middle panel**, Western blot showing the pattern of expression of activated ERK1/2 (pERK). **Bottom panel**, Quantification of Western blot data coming from 4 replicates. CNO induces a ∼76% increase of pERK1/2 in DRGs 2min after treatment of GFAP-hM3Dq mice as compared to WT mice (n=10 GFAP-hM3Dq mice, n=10 WT mice, Two-tailed unpaired t test: *P* = 0.0006). The level of pERK1/2 is not different from control levels 30min and 4h after treatment. Corresponding detailed quantification of the data is found in **Supplementary Table 3**.

**Supplementary Figure 5 (related to Fig. 1f).**
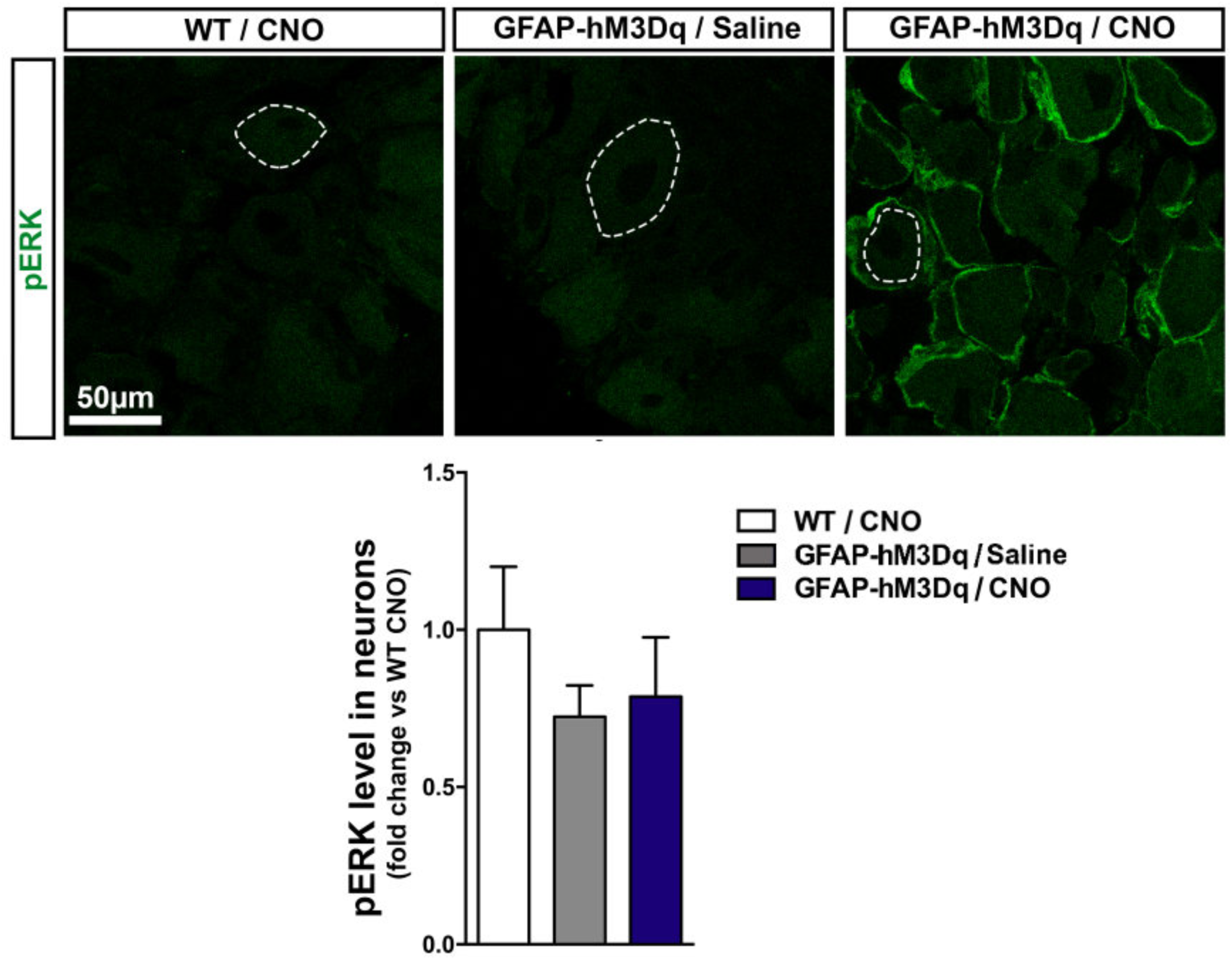
Immunofluorescence and confocal microscopy to study the expression of pERK in SCGs 2 min after 1 mg/kg CNO or saline i.p. treatment. Three groups of mice were used: CNO-treated WT mice, saline-treated-GFAP-hM3Dq mice and CNO-treated GFAP-hM3Dq mice. Mice were sacrificed 2min after treatment and their DRGs (L3, L4 and L5) were dissected, drop fixed in 4% paraformaldehyde and processed for immunohistochemistry. **Top panel**, confocal images showing pERK expression pattern. Note that pERK immunoreactivity is very low in SGCs from CNO-treated WT mice and saline-treated GFAP-hM3Dq mice as well in sensory neurons from the 3 groups of mice, contrary to SGCs from CNO-treatedGFAP-hM3Dq mice in which pERK immunoreactivity is high (green circles surrounding neuron soma, right). **Bottom panel**, Quantification of the data in large (∼40-50 μm diameter) sensory neurons (as delineated by dashed lines in top panel). CNO treatment does not induces an increase of pERK1/2 in sensory neuron soma 2min after CNO treatment in GFAP-hM3Dq mice as compared to saline-treated GFAP-hM3Dq and CNO-treated WT mice (n=4 CNO-treated GFAP-hM3Dq mice, n=4 saline-treated GFAP-hM3Dq mice, n=4 CNO-treated WT mice; Kruskal Wallis test: *P* = 0.5836). Corresponding detailed quantification of cellular characterization is found in **Supplementary Table 3**.

**Supplementary Figure 6 (related to Fig. 2a).**
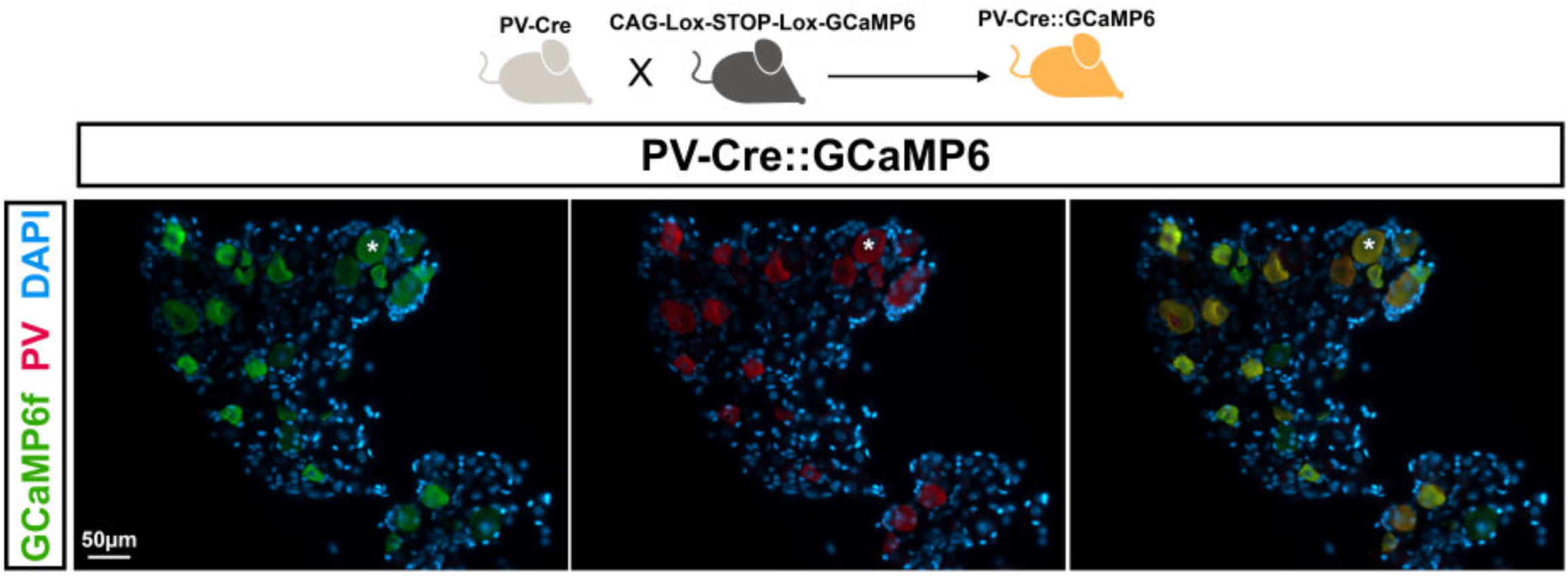
**Characterization of the PV-Cre::GCaMP6 double transgenic mouse line** showing that GCaMP6f protein (green) is expressed in ∼97% of parvalbumine (PV)-expressing proprioceptors (red) in DRGs (n=3 mice, n=6 slices 2 slices/mouse, n=37 PV-expressing proprioceptors). Cell nuclei are stained with DAPI (blue). One neuronal cell body is marked with asterisks. Images were acquired using an epifluorescence microscope. Corresponding detailed quantification of cellular characterization is found in **Supplementary Table 1**.

**Supplementary Figure 7 (related to Fig. 2a, c).**
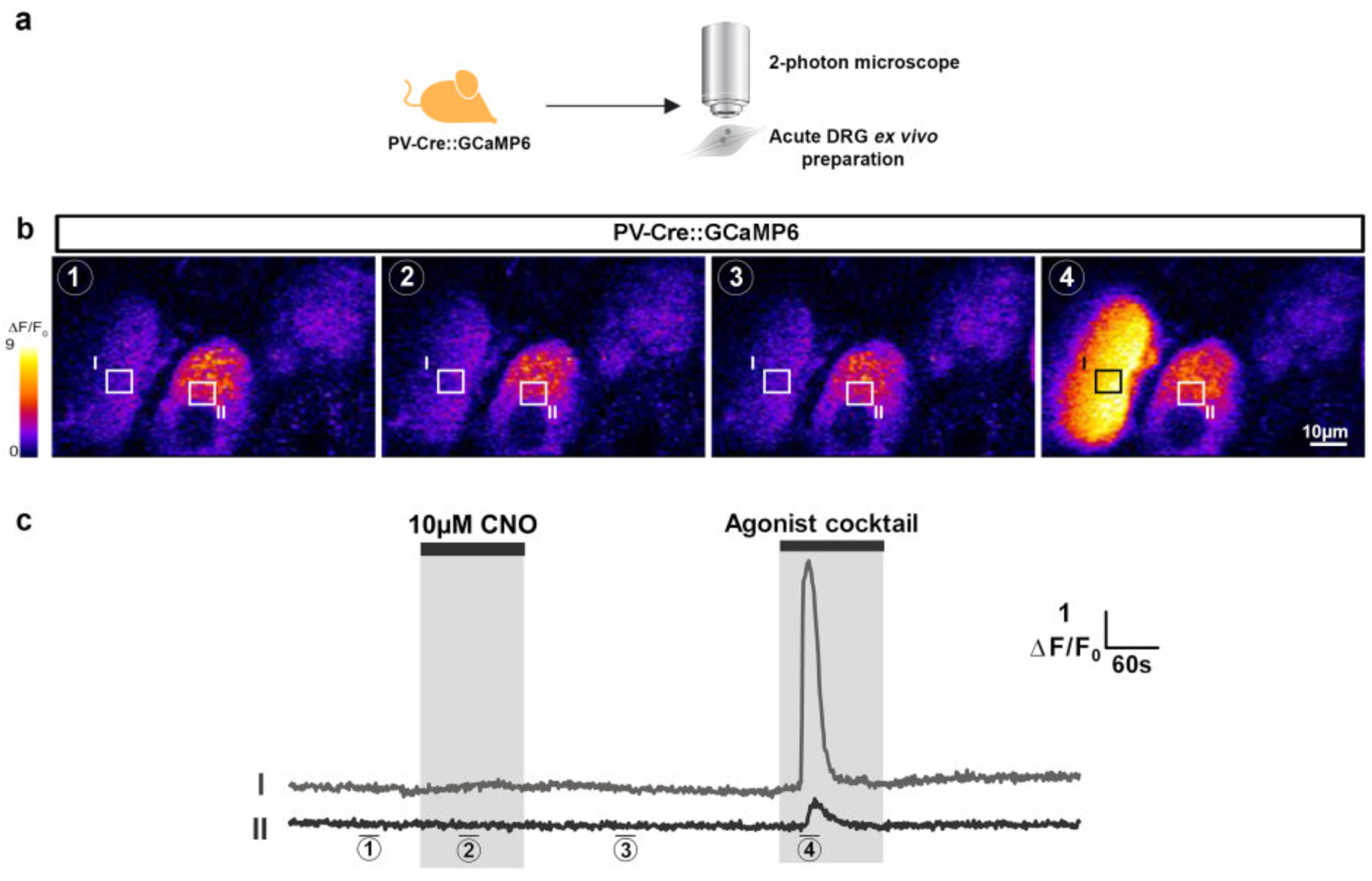
Inertness of CNO on proprioceptor activity. **a**, Experimental design. Two-photon Ca^2+^ imaging of *ex vivo* intact DRGs from PV-Cre::GCaMP6 double transgenic mice. **b**, Proprioceptor expressing GCaMP6f, outlined areas of interest (I, II) and Ca^2+^ increases during baseline ①, CNO application ②, wash ③ and cocktail application ④. **c**, Time course of Ca^2+^ traces in proprioceptors (I, II). A cocktail of ligands to endogenous G_q_ GPCR (1µM adenosine, 50µM ATP-γS, 10µM carbachol, 50µM DHPG, 200µM glutamate, 10µM histamine) has been applied after CNO washing to ensure proprioceptor viability. Corresponding detailed quantification of the data is found in **Supplementary Table 2**.

**Supplementary Figure 8 (related to Fig. 3a-d).**
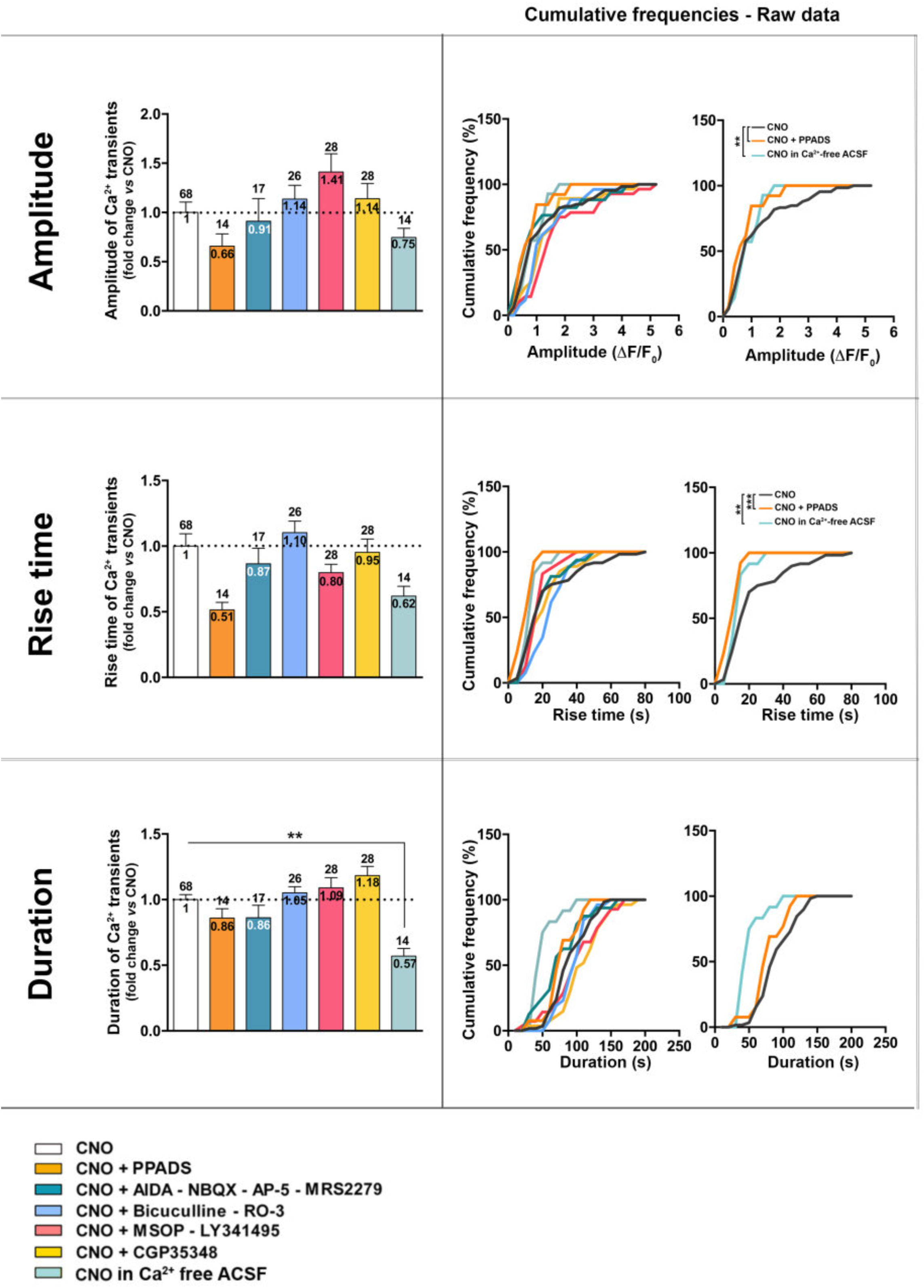
Pharmacological characterization of the receptors involved in CNO/hM3Dq/SGC-induced Ca^2+^ elevations in proprioceptors. Relative amplitude (**top panel**), rise time (**middle panel**) and duration (**bottom panel**) of the proprioceptor Ca^2+^ elevations in presence of CNO only or CNO combined with various transmitter receptor inhibitors alone or mixed: (1) 10 μM CNO (white), (2) 10 μM CNO + 100 μM PPADS (broad spectrum blocker of P2XRs & P2YRs) (orange), (3) 10 μM CNO + 100 μM AIDA (blocker of mGluR I) + 10 μM NBQX (blocker of AMPAR) + 50 μM AP5 (blocker of NMDAR) + 0.5 μM MRS2279 (blocker of P2Y_1_R) (dark blue); (4) 10 μM CNO + 50 μM Bicuculline (blocker of GABA_A_R) + 10 μM RO-3 (blocker of P2X_3_R) (blue); (5) 10 μM CNO + 100 μM MSOP (blocker of mGluR II) + 1 μM LY341495 (blocker of mGluR III) (red); (6) 10 μM CNO + 100 μM CGP35348 (blocker of GABA_B_R) (yellow); and (7) 10 μM CNO applied in Ca^2+^ free extracellular solution (light blue). **Left column**, histograms showing the quantification of the data normalized to CNO condition. **Right columns,** cumulative frequency distribution of proprioceptor response amplitude, rise time and duration (raw data), showing that the presence of PPADS or the absence of extracellular Ca^2+^ leads to decreases in the amplitude, rise-time and duration of the proprioceptor Ca^2+^ transients. Kruskal-Wallis and Kolmogorov-Smirnov tests were used: \*\**P* < 0.01, \*\*\**P* < 0.001; error bars indicate mean ± SEM. Corresponding detailed quantification and statistics are found in **Supplementary Table 5**.

**Supplementary Figure 9 (related to Fig. 4f).**
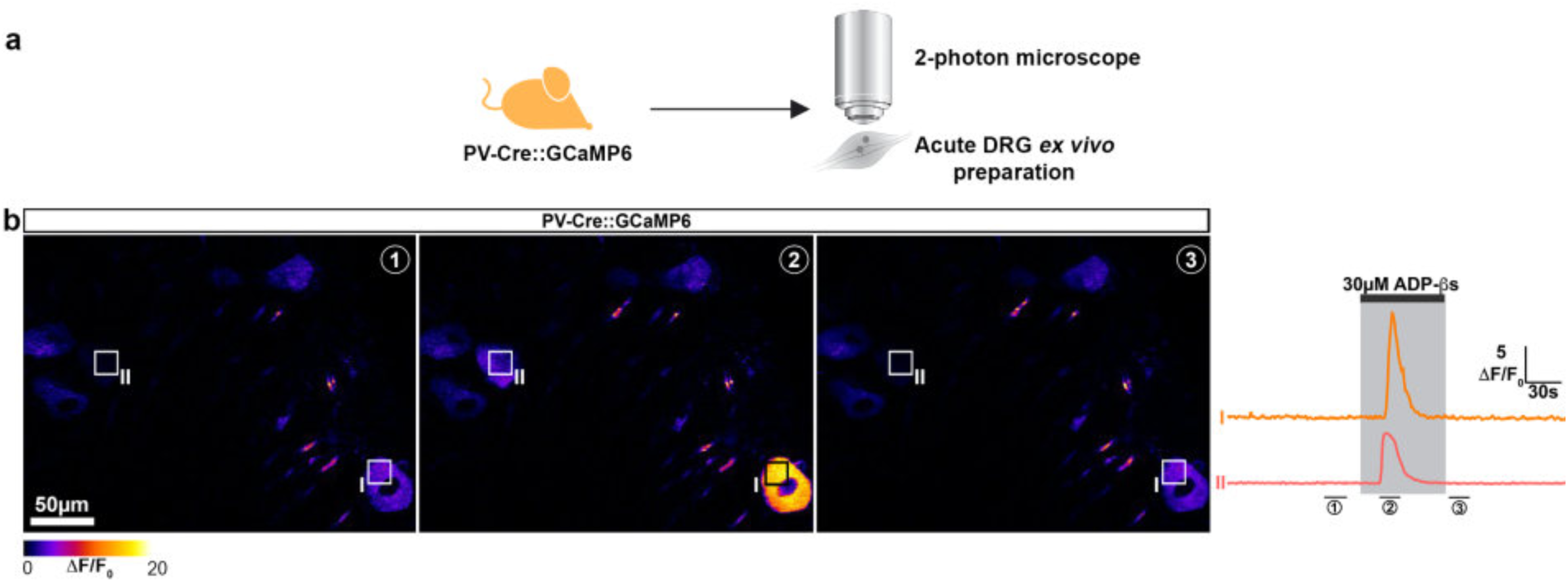
Proprioceptors express functional ADP receptors at their soma level. **a**, Experimental design. Two-photon Ca^2+^ imaging of *ex vivo* DRGs from PV-Cre::GCaMP6 double transgenic mice. **b**, **Left panel**, Cells expressing GCaMP6f, outlined areas of interest (I, II) and Ca^2+^ increases during baseline ①, ADP-βS application ② and wash ③. **Right panel** showing time course of Ca^2+^ traces in proprioceptors (I, II). Corresponding detailed quantification of the data is found in **Table 6**.

**Supplementary Figure 10 (related to Figs. 2a-d and 4a-e).**
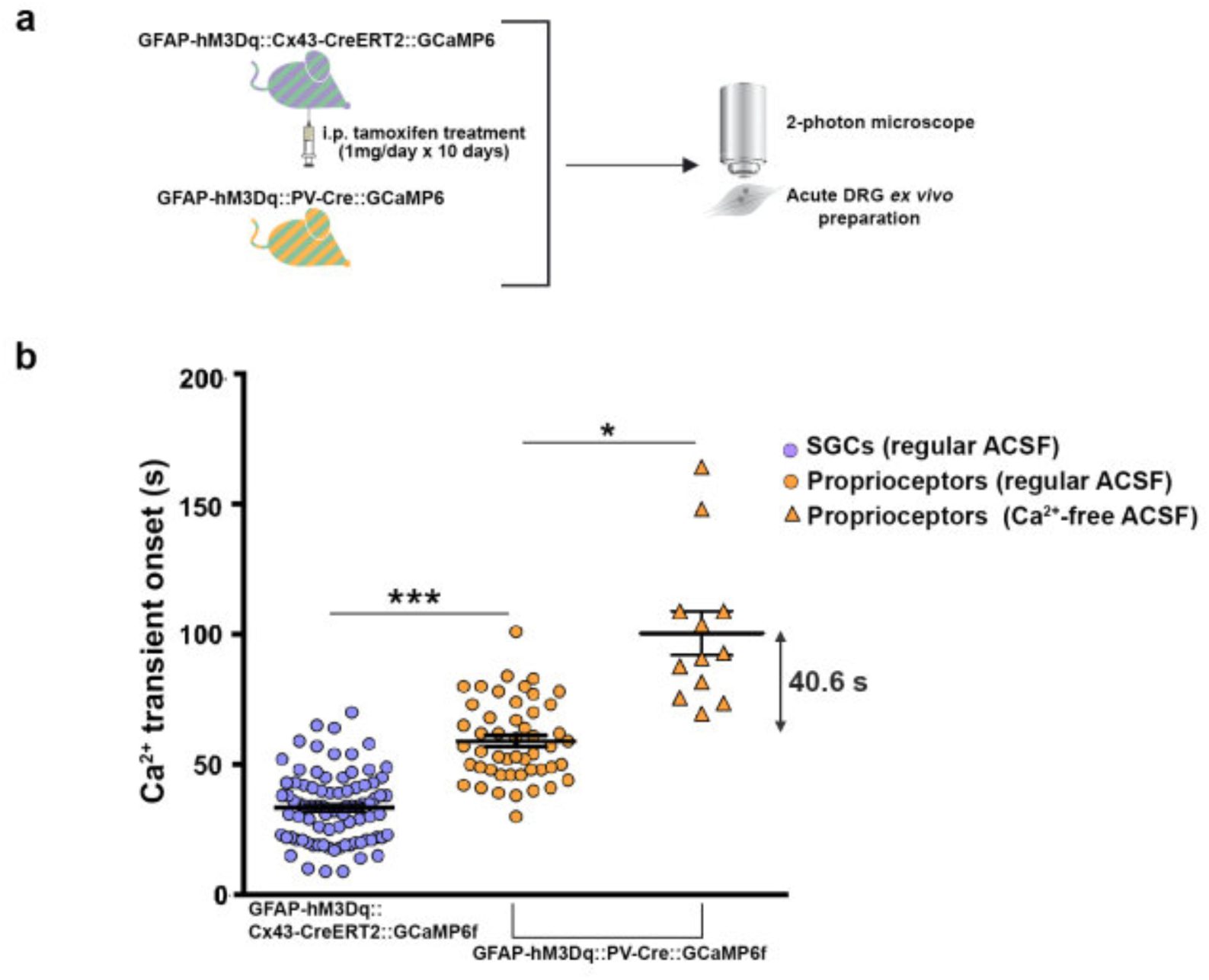
CNO/hM3Dq/SGC-induced Ca^2+^ responses in proprioceptors are delayed in absence of Ca^2+^ in the extracellular artificial cerebral fluid solution (ACSF). **a**, Experimental design. Two-photon Ca^2+^ imaging of *ex vivo* DRGs from GFAP-hM3Dq::Cx43-CreERT2::GCaMP6 and GFAP-hM3Dq::PV-Cre::GCaMP6 transgenic mouse lines expressing hM3Dq in SGCs and GCaMP6f either in SGCs or in proprioceptors, respectively. **b**, quantification of the data showing that Ca^2+^ elevations in SCGs (light blue circles, GFAP-hM3Dq::Cx43-CreERT2::GCaMP6 mice) occurs first, ∼25s before the induction of Ca^2+^ responses in proprioceptors (orange circles, GFAP-hM3Dq::PV-Cre::GCaMP6 mice) when CNO is applied in Ca^2+^-containing ACSF (see Fig. 2d for details). However, the onset of proprioceptor Ca^2+^ responses is delayed by ∼41 s (orange triangles, GFAP-hM3Dq::PV-Cre::GCaMP6 mice mice) when 10 μM CNO is applied in Ca^2+^ free ACSF as compared to the onset of the proprioceptor responses when CNO is applied in Ca^2+^-containing ACSF (Ca^2+^-containing ACSF: n=49 proprioceptors, n=23 DRGs, n=14 GFAP-hM3Dq::PV-Cre::GCaMP6 mice; Ca^2+^ free ACSF: =14 proprioceptors, n=15 DRGs, n=6 GFAP-hM3Dq::PV-Cre::GCaMP6 mice; Kruskal-Wallis test followed by Dunn’s multiple comparison test: *P* < 0.0001). Corresponding detailed quantification and statistics are found in **Supplementary Table 4**.

**Supplementary Figure 11 (related to Figs. 1-4 and Material & Methods).**
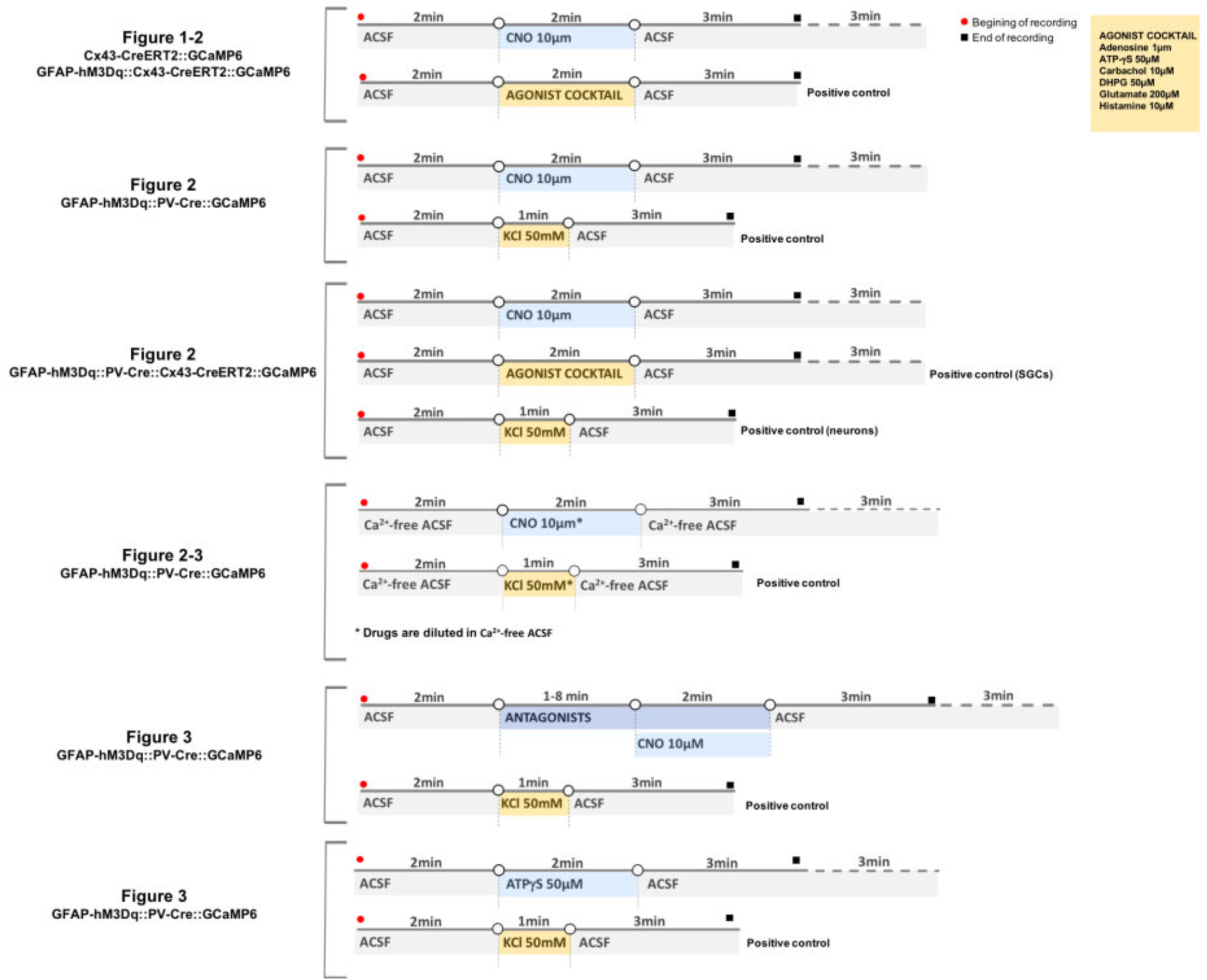
Designs of the different Ca^2+^ imaging experiments reported in the main manuscript and supplementary information document.

**Supplementary Table 1.** Detailed data and statistics relative to cellular characterization of GFAP-hM3Dq and PV-Cre::GCaMP6 mouse lines (immunohistochemistry experiments related to Fig. 1, Fig. 2 and **Supplementary Fig. 6**).

|  | n=mice | n=slice | n=neurons | n=SGCs (circle) | n= HA-positive neurons | % of HA-positive neurons | n= HA-positive SGCs | % of HA-positive SGCs |
| --- | --- | --- | --- | --- | --- | --- | --- | --- |
| GFAP-hM3Dq | 3 | 12 | 278 | 208 | 0 | 0 | 184 | 88,5% |

|  | n=mice | n=slice | n = PV-positive neurons | n = G6-positive PV-positive neurons | % of G6-positive PV-positive neurons |
| --- | --- | --- | --- | --- | --- |
| PV-Cre::GCaMP6 | 3 | 6 | 37 | 36 | 97,3% |

**Supplementary Table 2.**
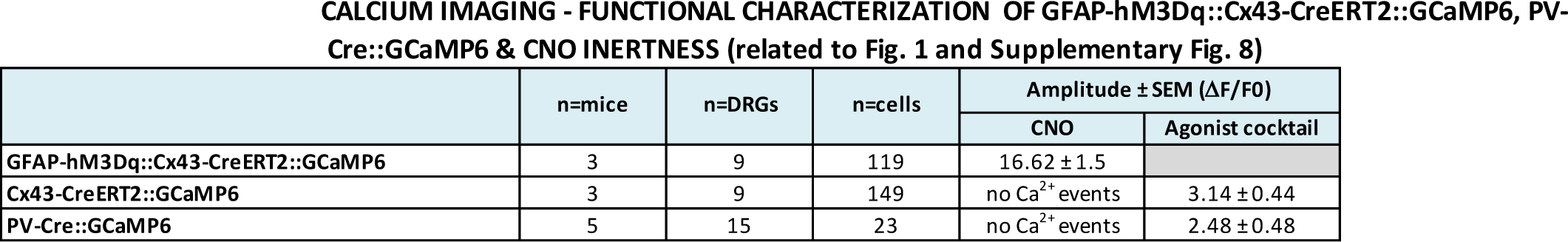
Detailed data and statistics relative to functional characterization of GFAP-hM3Dq::Cx43CreERT2::GCaMP6, Cx43-CreERT2::GCaMP6 and PV-Cre::GCaMP6 mouse lines (2-photon Ca^2+^ imaging experiments related to Fig. 1, Fig. 2 and **Supplementary Fig. 8**).

**Supplementary Table 3.**
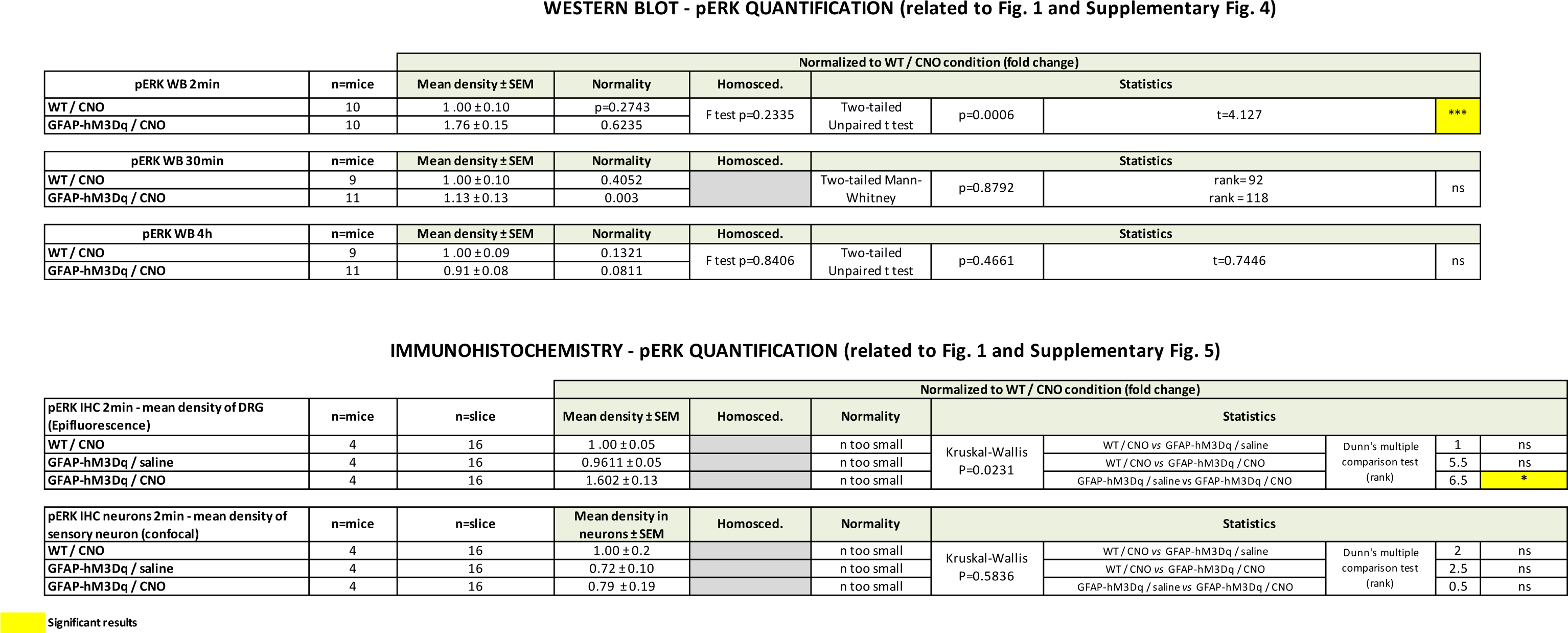
Detailed data and statistics relative to functional characterization of GFAP-hM3Dq mouse line (Western blot and immunohistochemistry experiments to reveal pERK expression levels; related to Fig. 1 and **Supplementary Fig. 4**).

**Supplementary Table 4.**
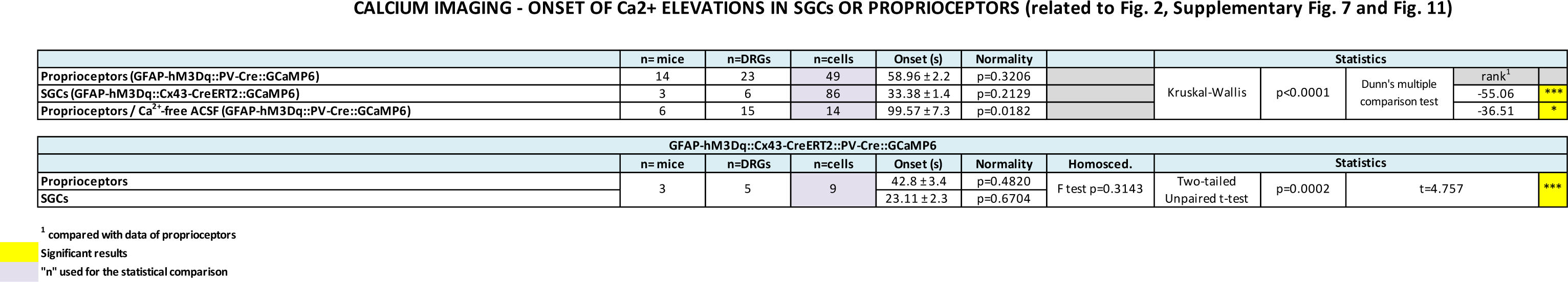
Detailed data and statistics relative to the discovery that the onset of CNO/hM3Dq-induced Ca^2+^ elevations in SGCs occur invariably ∼20-25 s before proprioceptor Ca^2+^ responses using GFAP-hM3Dq::Cx43-CreERT2::GCaMP6, GFAP-hM3Dq::PV-Cre::GCaMP6, and GFAP-hM3Dq::Cx43-CreERT2::PV-Cre::GCaMP6 transgenic mouse lines (2-photon Ca^2+^ imaging experiments related to **Fig. 2, Fig. 3** and **Supplementary Fig. 8** & **Fig. 10).**

**Supplementary Table 5.**
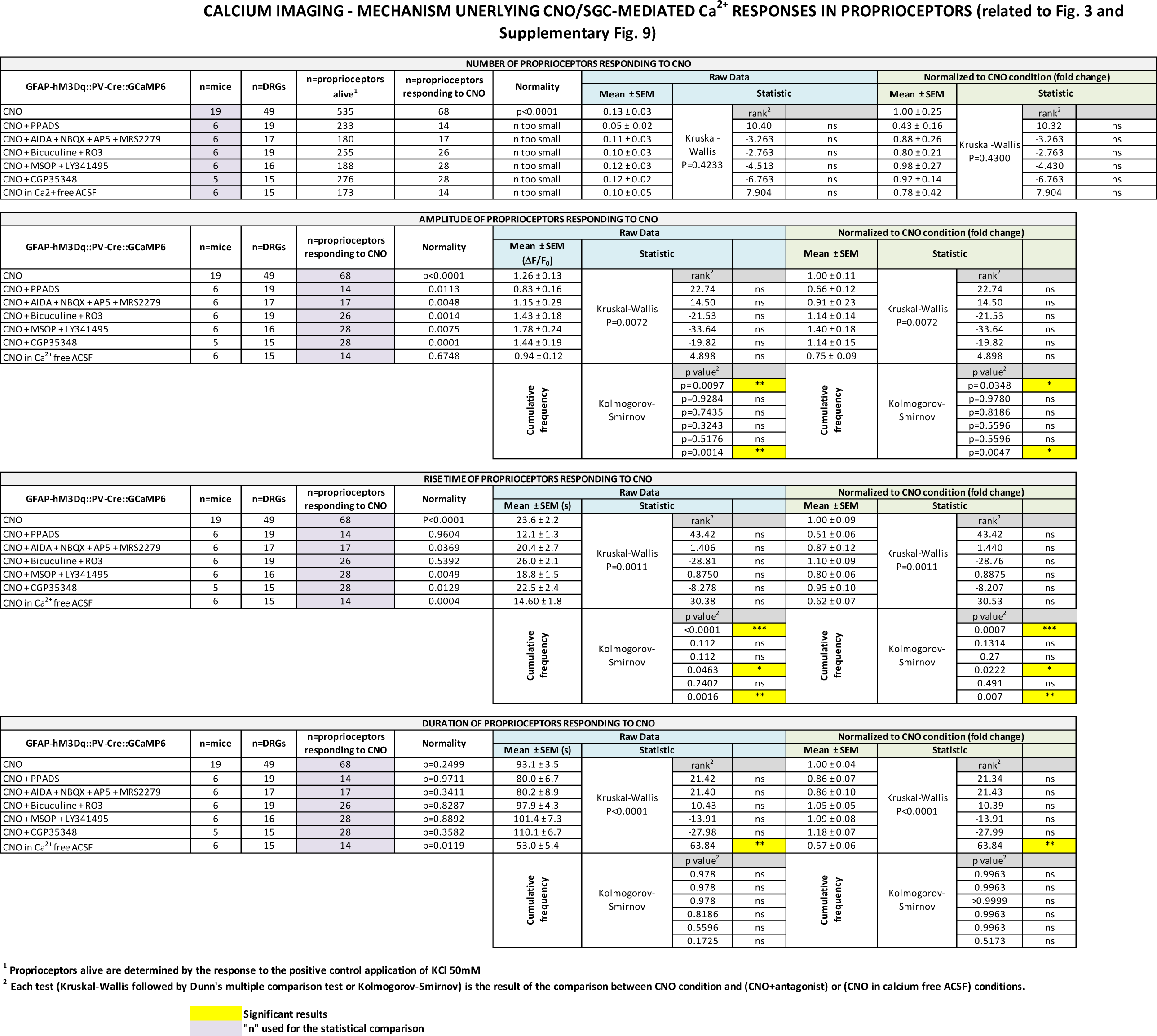
Detailed data and statistics relative to the discovery that purinergic P2XR and P2YR underlie CNO/SGC-mediated proprioceptor Ca^2+^ responses using GFAP-hM3Dq::PV-Cre::GCaMP6 mouse line (2-photon Ca^2+^ imaging experiments related to Fig. 4 and **Supplementary Fig. 8).**

**Supplementary Table 6.**
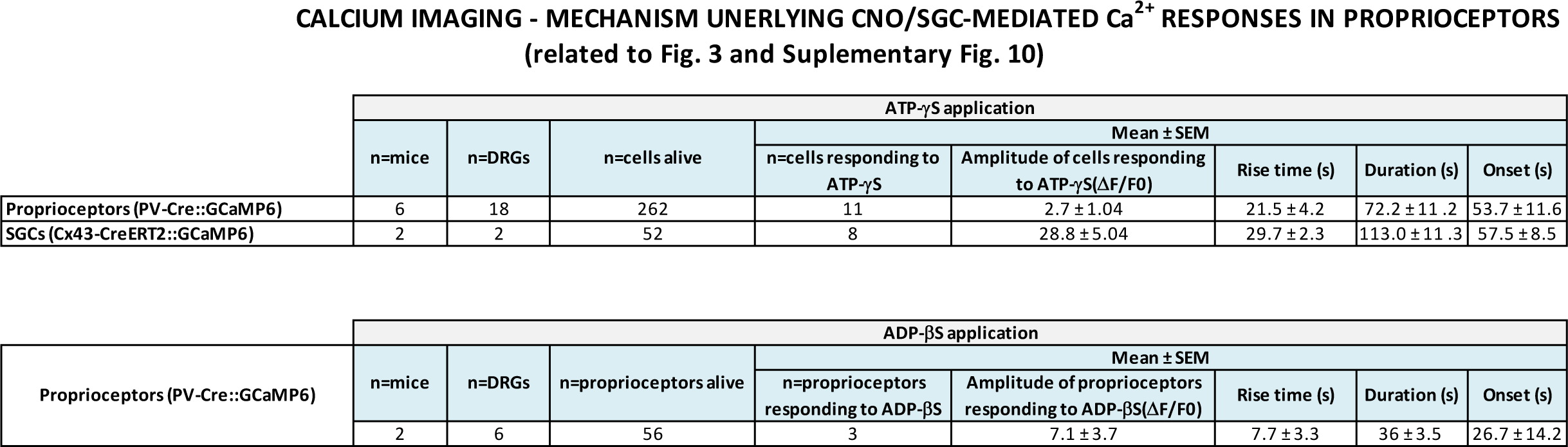
Detailed data and statistics relative to the discovery that ATPγS and ADPβS induce Ca^2+^ increases in proprioceptor using PV-Cre::GCaMP6 mouse line (2-photon Ca^2+^ imaging experiments related to Fig. 4 and **Supplementary Fig. 9).**

**Supplementary Table 7.**
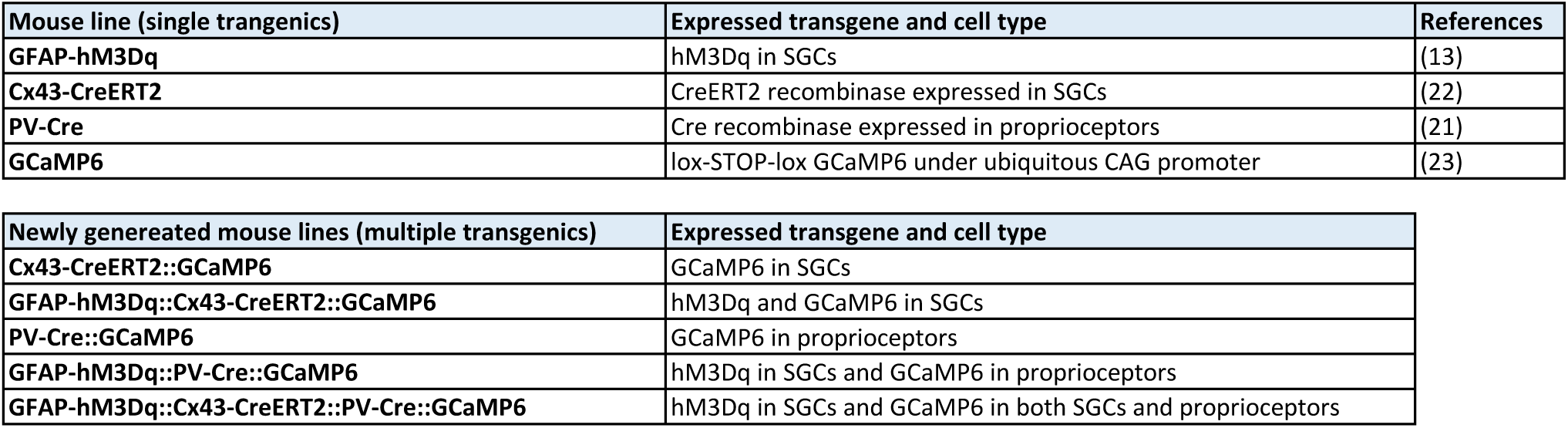
List of single, double, triple and quadruple transgenic mouse lines used in the study.

**Supplementary Table 8.** List of antibodies, drugs and reagents as well as corresponding concentrations used in the study. Abbreviations: IHC, immunohistochemistry; WB, Western blot.

| Antibodies |  |  |  |
| --- | --- | --- | --- |
| Name | Company/Cat. # | Species | Dilution |
| Primary antibodies |  |  |  |
| GAPDH | Milipore/MAB360 | Mouse | 1:20,000 (WB) |
| GFP | Invitrogen/A10262 | Chicken | 1:1,000 (IHC) |
| GLAST | Frontier Institute/Af660 | Rabbit | 1:5,000 (IHC) |
| HA tag | Roche/11867423001 | Rat | 1:1,000 (IHC) |
| Na<sup>+</sup>/K<sup>+</sup>-ATPase α-AF488 | Santa Cruz/sc-48345 AF488 | Mouse | 1:1,000 (IHC) |
| Neurofilament | Millipore/AB9568 | Rabbit | 1:1,000 (IHC) |
| pERK1/2 | Cell Signaling/4370 | Rabbit | 1:1,000 (WB & IHC) |
| Parvalbumin | Swant/PV27 | Rabbit | 1:1,000 (IHC) |
| Secondary antibodies |  |  |  |
| Alexa Fluor® 546 anti-rabbit | Invitrogen/A-11035 | Goat | 1:1,000 (IHC) |
| Alexa Fluor® 488 anti-chicken | Invitrogen/A-11039 | Goat | 1:1,000 (IHC) |
| HRP anti-rabbit | Jackson ImmunoResearch/111-035-003 | Goat | 1:10,000 (WB) |
| HRP anti-mouse | Jackson ImmunoResearch/115-035-003 | Goat | 1:10,000 (WB) |
| Drugs |  |  |  |
| Name | Company/Cat.# | Total bath application duration | Final concentration |
| Agonists – *ex vivo* |  |  |  |
| Adenosine | Abcam/ab120498 | 2min | 1μM (Ca<sup>2+</sup> imaging) |
| ADP-βS | Sigma/A8016 | 2min | 30μM (Ca<sup>2+</sup> imaging) |
| ATP-γS | Tocris/4080 | 2min | 50μM (Ca<sup>2+</sup> imaging) |
| Carbachol | Abcam/ab141354 | 2min | 10μM (Ca<sup>2+</sup> imaging) |
| CNO | NIH (Bryan Roth Lab) | 2min | 10μM (Ca<sup>2+</sup> imaging) |
| DHPG | Abcam/ab120020 | 2min | 50μM (Ca<sup>2+</sup> imaging) |
| Glutamate | Abcam/ab120049 | 2min | 200μM (Ca<sup>2+</sup> imaging) |
| Histamine | Sigma/H7125 | 2min | 10μM (Ca<sup>2+</sup> imaging) |
| Antagonists – *ex vivo* |  |  |  |
| AIDA | Enzo/ALX550101M025 | 3min | 100μM (Ca<sup>2+</sup> imaging) |
| AP5 | Abcam/ab120003 | 3min | 50μM (Ca<sup>2+</sup> imaging) |
| Bicuculine | Sigma/14340 | 3min | 50μM (Ca<sup>2+</sup> imaging) |
| CGP35348 | Abcam/ab120167 | 10min | 100μM (Ca<sup>2+</sup> imaging) |
| LY341495 | Abcam/ab120818 | 10min | 1μM (Ca<sup>2+</sup> imaging) |
| MRS 2279 | Tocris/2158 | 3min | 500nM (Ca<sup>2+</sup> imaging) |
| MSOP | Abcam/ab141381 | 10min | 100μM (Ca<sup>2+</sup> imaging) |
| NBQX | Abcam/ab120046 | 3min | 10μM (Ca<sup>2+</sup> imaging) |
| PPADS | Abcam/ab120009 | 3min | 100μM (Ca<sup>2+</sup> imaging) |
| R0-3 | Tocris/3052 | 3min | 10μM (Ca<sup>2+</sup> imaging) |
| Drugs – *in vivo* |  |  |  |
| Name | Company/Cat.# | Final concentration |  |
| CNO | NIH (Bryan Roth Lab) | 1mg/kg |  |
| Tamoxifen | Sigma/T5648 | 1mg/day (10mg/mL) |  |

**Supplementary Movie 1** (related to Fig. 1a and **Supplementary Fig.1)**

360° rotation of z-stack acquisition corresponding to **Supplementary Fig. 1.** HA in red, Na/K-ATPase in green.

**Supplementary Movie 2** (related to Fig. 2)

10 μM CNO application to *ex vivo* DRG from GFAP-hM3Dq::PV-Cre::GCaMP6 triple transgenic mice leads to Ca^2+^ responses in proprioceptors. CNO is applied at frame 120 until frame 240 (2-photon data acquisition rate: 1frame/s). Movie speed: 100 frames per second. AVI files were made with ImageJ.

10 μM CNO application to *ex vivo* DRG from GFAP-hM3Dq::Cx43-CreERT2::GCaMP6 triple transgenic mice induces Ca^2+^ elevations in SGCs. CNO is applied at frame 120 until frame 240 (2-photon data acquisition rate: 1frame/s). Movie speed: 100 frames per second. AVI files were made with ImageJ.

**Supplementary Movie 4** (related to Fig. 3)

10 μM CNO application to DRG from GFAP-hM3Dq::Cx43-CreERT2::PV-Cre::GCaMP6 quadruple transgenic mice shows Ca^2+^ responses occurring first in SGCs and then in neighboring proprioceptors. CNO is applied at frame 120 until frame 240 (2-photon data acquisition rate: 1frame/s). Movie speed: 100 frames per second. AVI files were made with ImageJ.

**Supplementary Movie 5** (related to **Material & Methods** – Calcium imaging section)

50mM KCl application to DRG from GFAP-hM3Dq::PV-Cre::GCaMP6 triple transgenic mice was used systematically as a positive control to determine proprioceptor viability (i.e. proprioceptors responding to KCl are considered alive). Movie speed: 100 frames per second. AVI files were made with ImageJ.

